# Exopolysaccharides of *Lactobacillus crispatus* mediate key balancing interactions with the vaginal mucosa

**DOI:** 10.64898/2026.02.05.703754

**Authors:** Vanessa Croatti, Caroline Dricot, Tom Eilers, Jelle Dillen, Tim Van Rillaer, Eline Cauwenberghs, Ilke Van Tente, Sam Bakelants, Dieter Vandenheuvel, Camille Allonsius, Isabel Pintelon, Sofie Thys, Wendy Mensah, Marina Naldi, Peter A. Bron, Stijn Wittouck, Irina Spacova, Carola Parolin, Beatrice Vitali, Sarah Lebeer

## Abstract

*Lactobacillus crispatus* is a dominant member of the healthy vaginal microbiota, yet the mechanisms by which it modulates host immunity remain poorly defined, in part due to the lack of tractable *in vivo* models. Here, we integrate bacterial genetics, *in vitro* epithelial systems, human-derived data and proteomic approach (Olink) to uncover a critical role for *L. crispatus* exopolysaccharides (EPS) in shaping the bacteria-vagina interactions. Comparative genomics identified a conserved EPS biosynthetic locus, with the priming glycosyltransferase gene *epsE* emerging as a regulatory node, in line with its distinct expression in human vaginal samples. Functional disruption of *epsE* abrogated *L. crispatus* EPS production and revealed its role for immune modulation. In human vaginal epithelial monolayers, EPS presence enhanced immune-regulatory (LAP TGF-beta-1) and anti-inflammatory (CST5) responses, whereas its absence triggered elevated pro-inflammatory cytokines (IL-1β, IL-6, IL-8) and matrix metalloproteinase (MMP-10). In a 3D vaginal organotypic model, EPS increased chemokines (CXCL5, CXCL6) linked to immune surveillance and the presence of the markers was validated in vaginal samples of healthy volunteers. These findings position EPS as a key immunomodulatory structure of *L. crispatus*, advancing our mechanistic understanding of host-commensal interactions and informing microbiome-based strategies to promote vaginal health.

## Introduction

Human reproduction and the overall health and well-being of women are significantly influenced by the unique symbiotic relationship between lactobacilli and the vaginal environment ^1^. While the composition of the vaginal microbiome can vary due to various environmental and lifestyle factors ^1,2^, a consistent dominance of *Lactobacillus crispatus* has been strongly linked to favorable health outcomes in various regions across the world ^3–5^. Such dominance of *L. crispatus* was also confirmed in healthy women in Western-Europe (Belgium) thanks to a large-scale citizen science initiative termed Isala ^6^, reinforcing the species’ potential role in supporting vaginal health. Despite some insights from preclinical studies have shed light on the protective activities of *L. crispatus* ^7–9^, the precise genes and molecules responsible for these beneficial effects, as opposed to those in less protective or pathobiont taxa, remain largely unknown. Mechanistic research on the vaginal microbiome is hindered by the lack of suitable animal models, as the symbiotic relationship with lactobacilli is unique to the human vagina. However, this gap is now being addressed through integrated analyses of human samples and well-designed *in vitro* experiments, supported by a steadily expanding arsenal of experimental models and study set-ups ^1,10–12^.

Bacterial cell wall components, particularly exopolysaccharides (EPS), represent a promising group of mediators in bacteria-host interaction mechanisms. These surface-associated molecules, together with other glycoconjugates, help establish the initial recognition and contact with the host by forming a distinctive molecular "barcode" ^13,14^. In *Lactobacillus* species, EPS are long, often branched homo-or heteropolysaccharides made from various monosaccharides, synthesized by enzymes encoded in organized gene clusters, including the key priming glycosyltransferase *epsE* ^15–17^. However, considerable strain- and species-level variability exists for *Lactobacillaceae* with some genes absent, redundant or scattered across the genome ^16^. This genomic diversity, together with the lack of precise knowledge about the functions and specificities of EPS biosynthetic enzymes, makes it difficult to establish clear correlations between genes and EPS structure. This challenge is particularly evident in key vaginal species like *L. crispatus*, where the pathways of EPS biosynthesis, its composition, and functional roles remain largely uncharacterized.

Genome editing is a powerful strategy for linking EPS biosynthesis to its ecological function, enabling targeted studies of specific genes involved ^18^. By comparing wild-type and EPS-mutants, this approach provides valuable phenotypic, structural and functional insights while preserving EPS in its native conformation, avoiding isolation methods that can alter the molecule’s structure and function ^19^. Yet, genetic modification remains challenging for vaginal lactobacilli isolates, *L. crispatus* included, limiting EPS research primarily to gut-associated strains, such as *Lacticaseibacillus rhamnosus* GG ^17^, *Lacticaseibacillus casei* Shirota ^20^, *Lactiplantibacillus plantarum* WCFS1 ^21^ and *L. plantarum* Lp90 ^22^. These studies have highlighted the pivotal role of EPS in bacteria-gut interactions, including adhesion to the host and modulation of the immune system. While different strains of gut lactobacilli have been shown to interact with gut and immune cells via pattern recognition receptors (PRRs) such as Toll-like receptor 2 (TLR2), and C-type lectins such as DC-SIGN, and CD14^23^, it remains unclear whether EPS are directly recognized or merely mask other bacterial components that interact with these receptors. Moreover, the scope of the immune responses captured in many studies remains limited due to the narrow range of cytokines and biomarkers typically assessed using targeted qPCR or ELISA methods ^24,25^. However, recent advances in high-throughput immune profiling technologies, such as multiplex cytokine assays Luminex panel ^26,27^ or Olink panels (Olink, Uppsala, Sweden) and transcriptomic approaches like RNA sequencing^28^ are enabling more comprehensive and systematic analyses of host-microbiome interactions. These tools, when applied to *in vitro* coculture systems and in human samples, including analyses of clinical samples such as vaginal swabs^29^, offer new opportunities to unravel the complexity of immune modulation by lactobacilli.

In this study, we investigate the role of *L. crispatus-*derived EPS in maintaining vaginal host symbiosis. Leveraging a unique biorepository of vaginal samples, we combine comparative genomics of public and in-house strains, *in vivo* transcriptomic analysis of the *eps* gene expression exploiting publicly available datasets, targeted *epsE* mutagenesis, and phenotypic gene-function studies using epithelial monolayers and 3D biomimetic models, with validation of immune biomarkers expression in healthy volunteers. Through this integrated approach, we assess how EPS shapes bacteria–host interactions in the vaginal niche, including modulation of host immunity. Our findings provide new insights into the contribution of EPS to the symbiotic relationship between *L. crispatus* and the human vagina, advancing our understanding of the microbial mechanisms that underpin vaginal health.

## Results

### *L. crispatus* has an EPS biosynthetic cluster expressed in the human vagina

First, we aimed to map *L. crispatus* genetic potential for EPS production through a comparative genomics approach. To achieve this, the amino acid sequence of a priming glycosyltransferase protein (encoded by the gene *epsE*) from a *Lactobacillus gasseri* strain associated with EPS production^30^ was used as a query to identify the EPS gene cluster in 7 selected in-house *L. crispatus* strains isolated from vaginal swabs, chosen for their suitability for functional characterization, including their ability to grow under conditions used for preparing competent cells and their antibiotic resistance, used for gene replacement (**Supplementary Information, Tab. S1**). All in-house strains contained an EPS gene cluster (**Fig. 1A**), with the first five genes highly conserved: LytR transcriptional regulator (*epsA*, minimum 97% pairwise nucleotide sequence identity), tyrosine kinase modulator (*epsC*, 93%), tyrosine kinase (*epsD*, 95%) and phosphotyrosine phosphatase (*epsB*, 94%), involved in glycan chain length regulation, and the undecaprenyl-phosphate galactose phosphotransferase [EC 2.7.8.6 – KEGG Orthology (KO)] (priming glycosyltransferase- *epsE*, 87%). The remaining EPS genes, which encode other glycosyltransferases, flippases, a polymerase, and various accessory enzymes, were conserved to a lesser extent, indicating a potential variability in EPS structure and/or composition among the strains.

**Figure 1.**
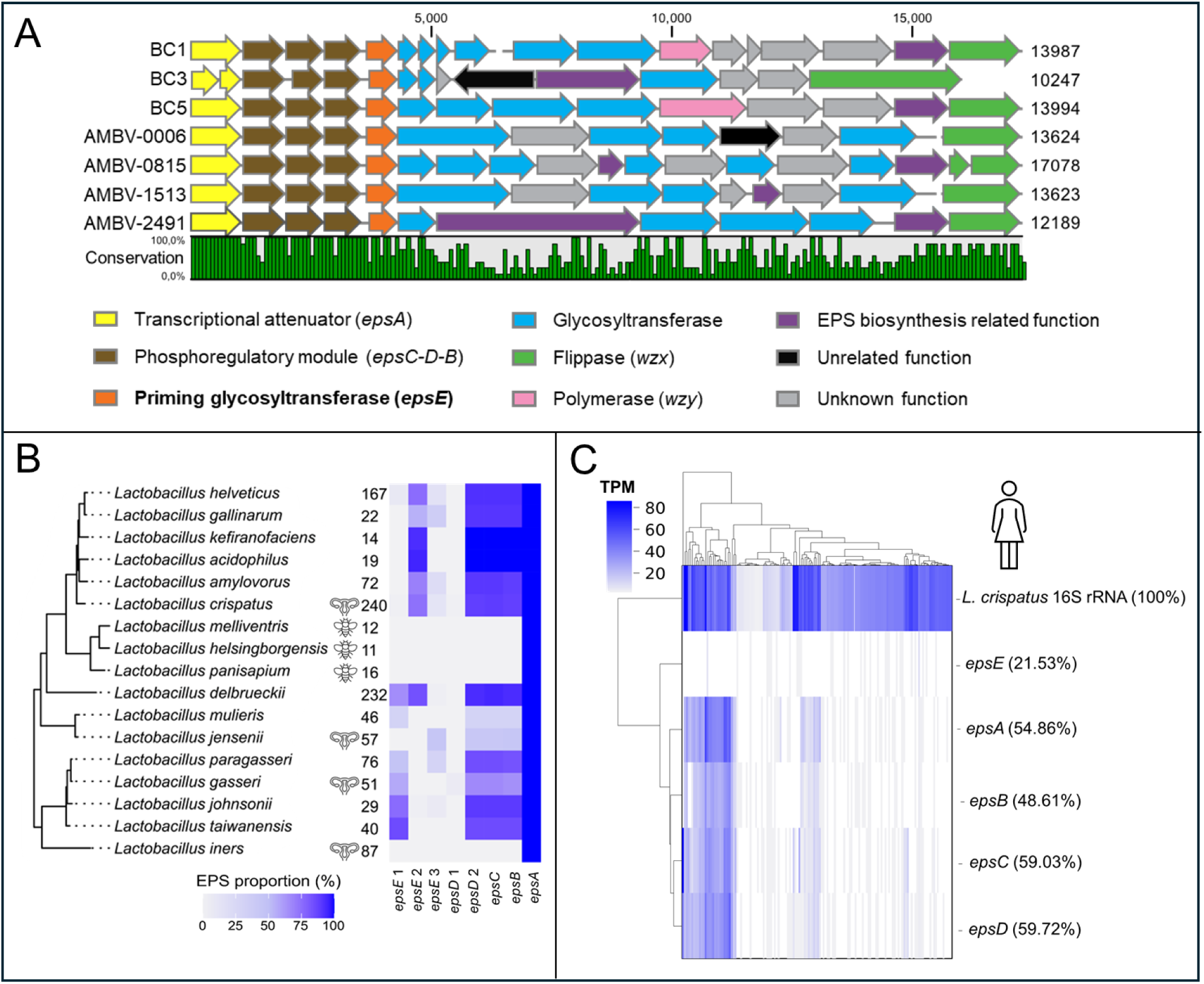
*L. crispatus* eps genes conserved features and their expression in the human vagina. (A) Schematic representation of the EPS gene cluster in selected *L. crispatus* strains and sequences alignment. Nucleotide sequence conservation between the strains is reported in the line plot (%). The legend of the annotated genes is shown. (B) Pangenome analysis on the *epsA, epsC, epsD, epsB, epsE* genes within the genus Lactobacillus and isolates from different habitats. The proportion of unique strains containing the orthogroup to which each gene is assigned is shown, along with the total number of unique strains. Species were only retained if they contained 10 unique strains. The female reproductive tract illustration highlights the four dominant vaginal *Lactobacillus* species*: L. crispatus, L. jensenii, L. gasseri,* and *L. iners*; the bee illustration shows the species (bee)-adapted. (C) Analysis of 144 vaginal sample transcriptomes for the expression of *epsA, epsC, epsD, epsB, epsE* genes of selected *L. crispatus*. To allow for comparison, L. crispatus 16S rRNA TPM values have been divided by 1000 due to their significantly higher expression (∼1000x), while TPM values for eps transcripts are presented at their original scale.

To explore the conservation of the EPS biosynthetic gene cluster within the genus *Lactobacillus* and a possible association with adaptation to the vagina versus other major habitats, we analyzed the presence of these five conserved *eps* genes in a dereplicated dataset of 1609 unique genomes (**Fig. 1B**). While pangenome analysis revealed that *epsA* was present in all members of the genus *Lactobacillus* studied, including the species associated with the vaginal community state types (*L. crispatus, L. gasseri, L. iners* and *L. jensenii*), its functional relevance is likely limited. In contrast, *epsE,* which encodes the priming glycosyltransferase responsible for the initiation of EPS biosynthesis, showed the highest variability and was distributed into different orthogroups, suggesting functional diversification. Of note, *epsE* was found to belong to distinct orthogroups across different vaginal species, suggesting species-specific adaptations to the vaginal niche. The structural genes *epsBCD* were highly prevalent in *L. crispatus* and *L. gasseri* but were only detected in a minority of *L. jensenii* analyzed. Notably, *L. iners* harbored only *epsA*, indicating that this species is likely unable to produce EPS. In addition, the closely related invertebrate (bee)-adapted clade containing taxa such as *L. melliventris, helsingborgensis* and *L. panisapium* have also lost the *eps* genes analyzed with exception of *epsA*.

To assess the expression of EPS by *L. crispatus* in humans, the transcriptional activity of the conserved *epsA*, *epsC*, *epsD*, *epsB*, and *epsE* genes was analyzed using publicly available RNA-seq data derived from vaginal microbial communities of 144 healthy women ^31^. Out of 144 samples, 114 expressed *L. crispatus* 16S rRNA, of which 105 had at least one *eps* transcript (**Fig. 1C**). The regulatory genes (*epsA*, *epsC*, *epsD*, or *epsB*) were detected in over 48% of the *L. crispatus* samples, indicating that these genes are commonly expressed by *L. crispatus* in the vagina.. In contrast, *epsE* was detected in fewer samples (21.5%), suggesting it may be more tightly regulated and serve as a key control point for EPS production.

### Deletion of *epsE* in vaginal *L. crispatus* strain impacts EPS production and composition

We aimed to investigate the functional role of cell wall EPS by generating loss-of-function mutants through targeted deletion of *epsE*. To knockout *epsE* in multiple strains, a semi-generic mutagenesis plasmid (pAMB5815) was designed using a subgroup of in-house strains with two highly conserved homologous regions (∼550 base pairs) flanking *epsE*, tolerating a maximum of 7% mismatches (BC1, BC5 and AMBV-0815) (**Supplementary Information, Fig. S1**). Among the tested strains, only *L. crispatus* AMBV-0815 exhibited a sufficiently high transformation efficiency and yielded a successful colony using pAMB5815 (**Supplementary Information, Tab. S2**), displaying the anticipated phenotype and genotype, checked with PCR and Sanger sequencing (**Supplementary Information, Fig. S2**). The mutant obtained, named *L. crispatus* AMBV-0815Δ*epsE* (EPS-mutant), was used for further phenotypic investigations.

Absence of *epsE* did not significantly affect the growth rate of the EPS-mutant compared to AMBV-0815 wild-type (0.30 ± 0.04 h^−1^ vs. 0.32 ± 0.04 h^−1^, respectively), which was important to exclude pleiotropic effects (**Supplementary Information, Fig. S3**). We then confirmed a reduction in total cell wall EPS in the EPS-mutant strain by different approaches (**Fig. 2**). Transmission electron microscopy (TEM) (**Fig. 2A**) showed that *L. crispatus* EPS-mutant lacked the extracellular polysaccharide layer observed in the wild-type (∼602 ± 151 nm of thickness, n=3) and displayed a cellular rough phenotype in contrast to the smooth cell morphology of the wild-type. Crude capsular sugars isolation showed a ∼4-fold decrease in yield (1.24 ± 0.59 μg/10⁹ CFU for the EPS-mutant vs. 5.04 ± 1.31 μg/10⁹ CFU for the wild-type) (**Fig. 2B**) and an alteration in the monosaccharide’s relative abundance (**Fig. 2C**). Capsular sugars from *L. crispatus* wild-type consisted of 9 different monosaccharides (**Supplementary Information, Fig. S4**): D-Galactosamine, D-Ribose, D-Glucosamine, D-Galactose, D-Glucose, D-Rhamnose, D-Xylose, D-Fucose, and D-Mannose, with D-Galactosamine, D-Ribose, D-Glucosamine and D-Galactose being the most abundant (≥ 85% of total sugars composition). The deletion of *epsE* led to a significant reduction (Student’s t-test with Welch correction, p<0.001, n=3) in D-Glucosamine (10% vs 16% relative abundance) and D-Galactose (7% vs 10% relative abundance) content, and, notably, an increase in D-Galactosamine (40% vs 31%), indicating not a full capsular sugar or cell wall glycoconjugate depletion, in line with the presence of other surface glycans than the EPS under study (**Fig. 2C**).

**Figure 2.**
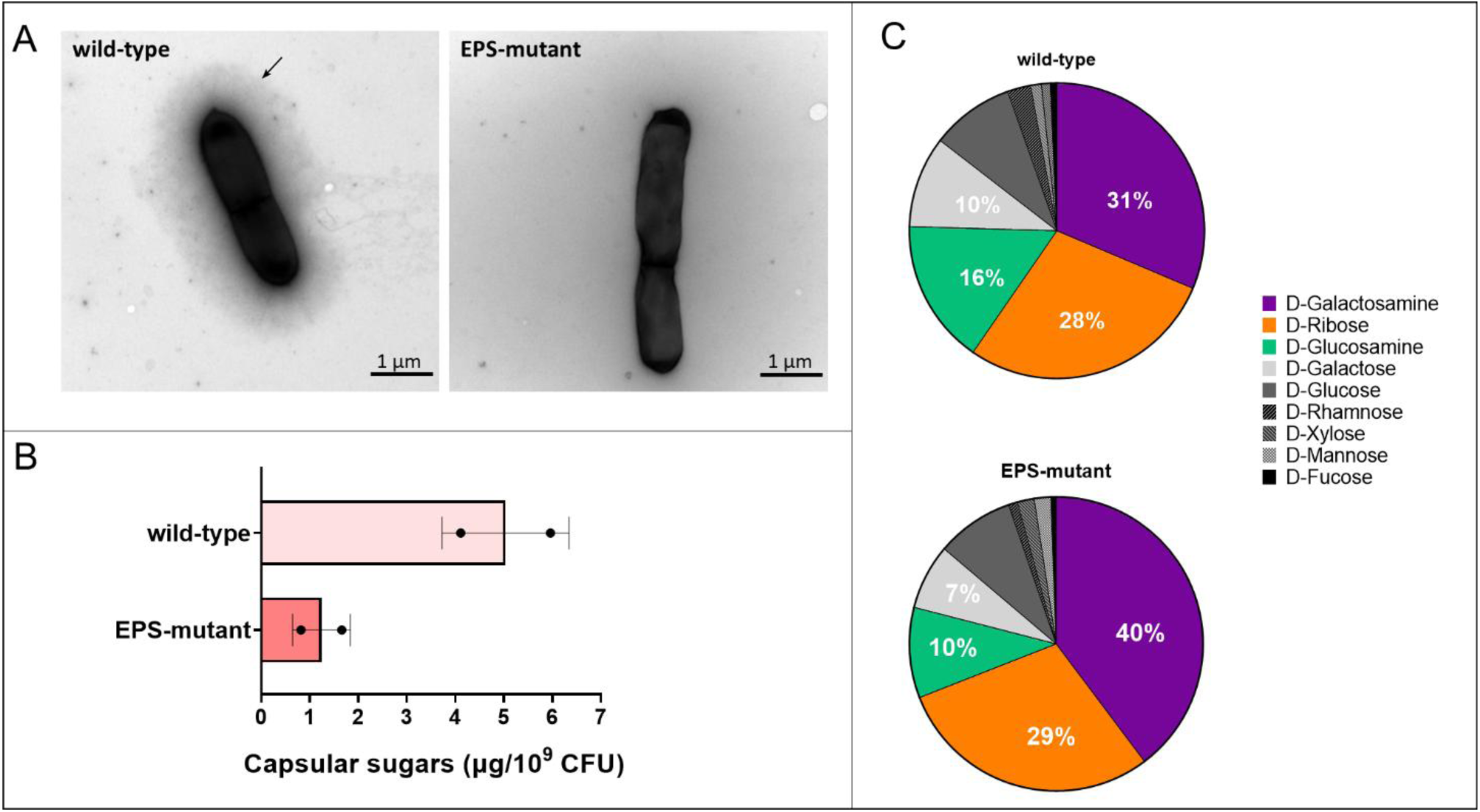
Impact of *epsE* deletion on capsular sugars yield and composition. (A) TEM micrographs of *L. crispatus* AMBV-0815 wild-type and EPS-mutant. Sugars layer indicated with black arrow. The scale bar is indicated. (B) Quantification of crude capsular sugars isolated from AMBV-0815 wild-type and EPS-mutant. Data is reported as mean ± SD (n = 2). (C) Relative abundance of monosaccharides (LC-MS) in capsular sugars recovered from AMBV-0815 wild-type and EPS-mutant (n=3).

### The presence of EPS affects the adhesive capacity of vaginal *L. crispatus*

Subsequently, we explored the role of *L. crispatus* cell wall EPS in bacteria biofilm formation and host-adhesion by comparing the adhesive capacities of *L. crispatus* AMBV-0815 wild-type and EPS-mutant. The EPS-mutant displayed a 2-fold reduction in biofilm formation compared to the wild-type strain (52% vs 100%), in line with a role for EPS in biofilm formation (**Fig. 3A**). However, a ca. 2-fold increase in adherence to vaginal epithelial cells (VK2/E6E7) monolayer was observed using direct (microscope counting) and indirect (plating out) approaches (**Fig. 3B**). Fluorescence microscope imaging of FITC-stained AMBV-0815 wild-type and EPS-mutant (**Supplementary Information**, **Fig. S5**) further confirmed the enhanced adhesive capacity of the EPS-mutant strain on the vaginal epithelium compared to the wild-type counterpart. These findings suggest that EPS plays a modulating role in *L. crispatus* AMBV-0815 interactions with abiotic and biotic surfaces, promoting biofilm formation, while reducing adhesion to vaginal epithelial cells. The absence of any direct and/or indirect cytotoxic effect of vaginal epithelial cells on bacterial viability, which could potentially influence adhesion rates, was verified and reported in **Supplementary Information Fig. S6**. For this reason, we concluded that the observed difference in adhesion is primarily attributed to altered adhesive capacity of the mutant strain, rather than to increased survival relative to the wild-type.

**Figure 3.**
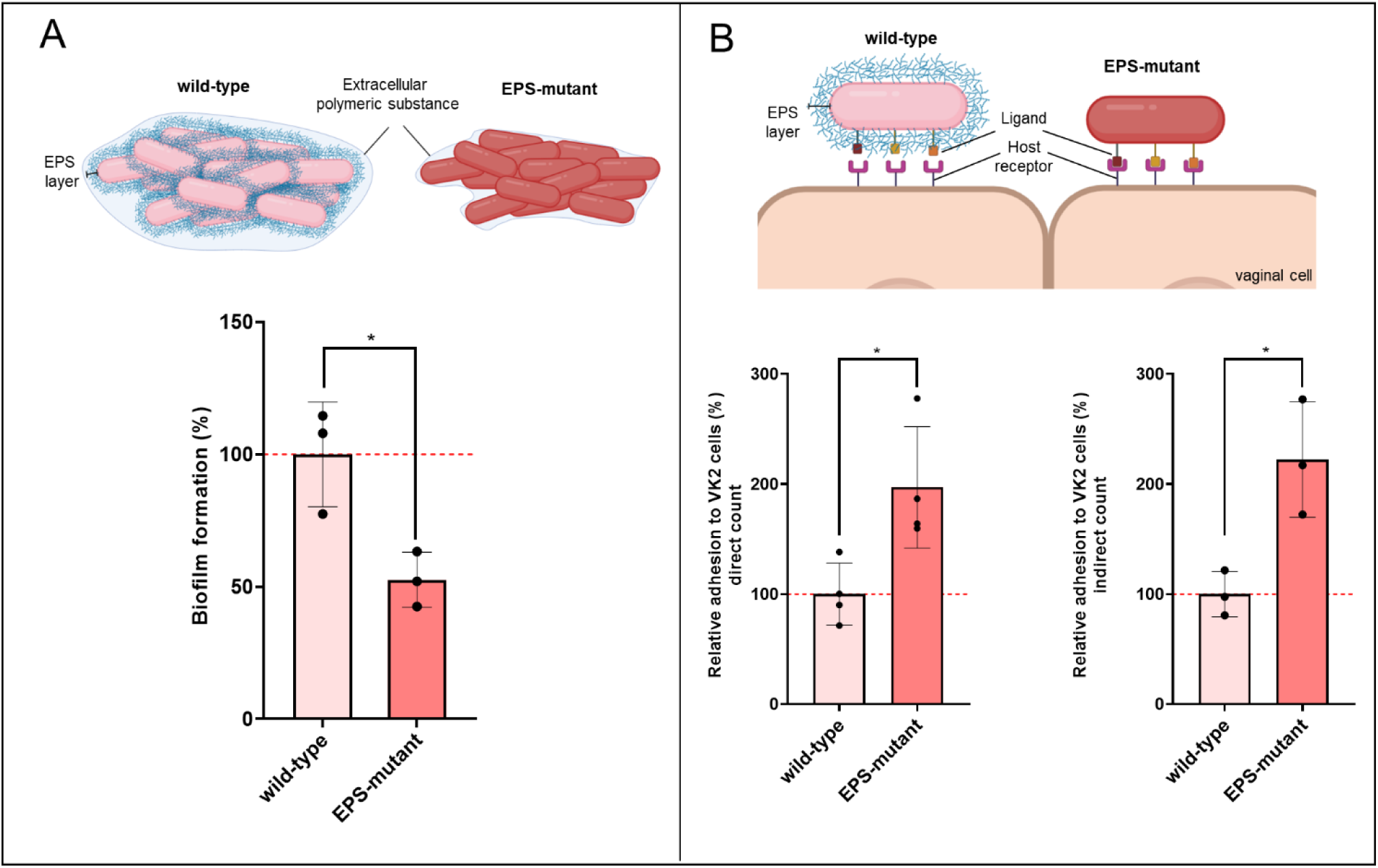
Role of EPS in *L. crispatus* biofilm formation and adhesion to immortalized vaginal monolayer. (A) Biofilm formation of *L. crispatus* AMBV-0815 wild-type and EPS-mutant on abiotic surfaces. The percentages of biofilm formation are reported in relation to AMBV-0815 wild-type biofilm values (set as 100%, red dotted line) (n=3). (B) Adhesion of AMBV-0815 wild-type and EPS-mutant on VK2/E6E7 cells monolayer evaluated by direct (plating out) and indirect (microscope counting after May-Grunwald/Giemsa staining) methods. The percentages of adhesion are reported in relation to AMBV-0815 wild-type adhesive values (set as 100%, red dotted line) (n=3-4). Data are reported as mean ± SD. Welch’s t-test (*p < 0.05). Figure made with *Biorender.com*

### The presence of EPS modulates *L. crispatus* interactions with the vaginal immune system

We then evaluated the role of *L. crispatus* cell wall EPS in the host innate immune responses by comparing immune parameters of different cell models exposed to the wild-type and EPS-mutant strains, allowing us to study EPS in its native conformation as presented to host cells. We first employed human reporter monocytes (THP1-Dual) to monitor *L. crispatus* AMBV-0815 wild-type and EPS-mutant interaction with key innate immune pathways involved in the production of cytokines, chemokines, and interferons as critical components of antibacterial and/or antiviral immune defense. Specifically, we examined the activation of the Nuclear Factor kappa-light-chain-enhancer of activated B cells (NF-κB) and Interferon Receptor Factors (IRF) transcription pathways. As shown in **Fig. 4A**, both *L. crispatus* AMBV-0815 wild-type and the EPS-mutant significantly activated NF-κB and IRF pathways compared to non-exposed cells after 24 hours. Notably, the EPS-mutant induced a stronger activation of IRF pathway compared to the wild-type, showing a ∼1.5-fold increase (11397 vs. 7707 luminescence, respectively, p = 0.002, n=3). While NF-κB pathway activation also increased with the EPS-mutant, the difference was not statistically significant under the tested conditions (One-way ANOVA with Tukey correction, p=0.25, n=3).

**Figure 4.**
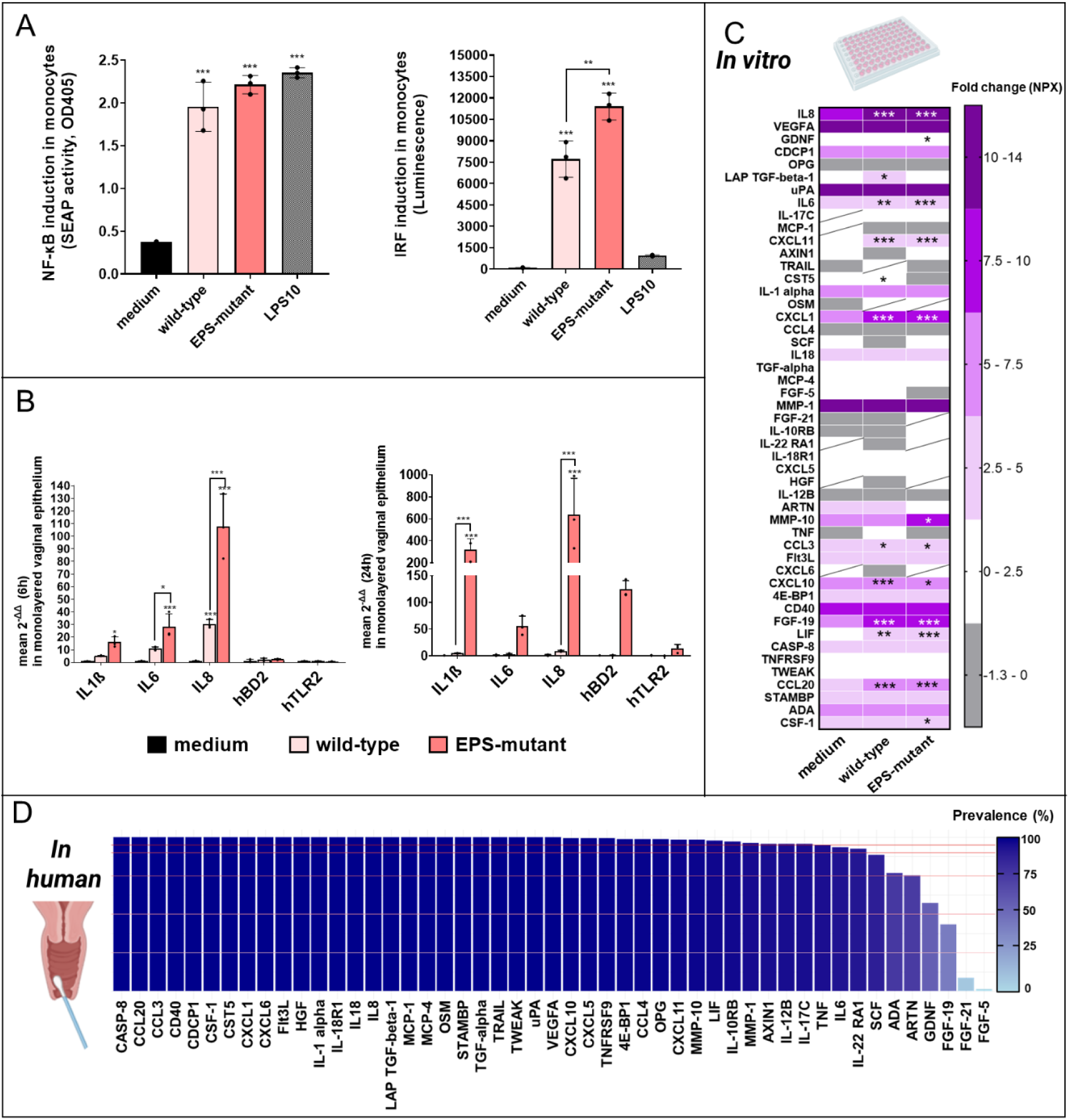
Role of EPS in *L. crispatus* interactions with the innate immune system. (A) *L. crispatus* AMBV-0815 wild-type and EPS-mutant stimulation of NF-κB and IRFs in human THP1-Dual monocytes after 24h exposure. RPMI medium was used as negative control. LPS at 10 ng/mL (LPS10) was used as control NF-κB and IRF inducer. Data are represented as mean ± SD (n=3). One-way ANOVA with Tukey correction. (B) Cytokine and other immune markers expression in immortalized vaginal epithelial cells (VK2/E6E7) (RT-q-PCR) after 6h and 24h exposure to the *L. crispatus* AMBV-0815 wild-type and EPS-mutant. KFSM medium was used as a negative control. Fold-change is expressed in respect to the negative control. Data are represented as mean ± SD (n=3). (C) Effect of *L. crispatus* AMBV-0815 wild-type and EPS-mutant on the inflammation panel in vaginal epithelial cells (VK2/E6E7) (24h) (Olink® assay). KSFM medium was used as negative control. Data are reported as fold-change (NPX) of 49 inflammatory biomarkers. Biomarkers with more than 55% samples below the limit of detection (LOD) are represented by cells with a line. Statistical comparisons relative to the negative control are presented in the heat map (n=3). Two-way ANOVA with Tukey correction (*p = 0.033, **p = 0.002, ***p < 0.001). (D) Prevalence (%) of 49 immune biomarkers detected above LOD in human vaginal samples of healthy volunteers, as identified by the Olink® assay in monolayer vaginal epithelium. The horizontal lines indicate a 95%, 90%, 75%, 50% and 25% prevalence. Figure made with *Biorender.com*

Next, we aimed to investigate how the presence of EPS on *L. crispatus* modulates the immune response of vaginal mucosal cells to bacteria. A monolayer of the human vaginal epithelial cells (VK2/E6E7) was exposed to *L. crispatus* AMBV-0815 wild-type and EPS-mutant strains for 6 and 24 hours and the expression levels of molecules involved in general host-microbe interactions (human toll-like receptors 2 – hTLR2), inflammation (human pro-inflammatory cytokines IL1β, IL6, IL8) and antimicrobial responses (human beta-defensin-2 - hBD2) were then established using reverse transcription qPCR. As shown in **Fig. 4B**, exposure to *L. crispatus* AMBV-0815 wild-type at 6 and 24 hours did not induce significant changes in the transcription of these immune markers in vaginal cells, except for an increase in IL8 levels at 6 hours. In contrast, the EPS-mutant upregulated IL1β, IL6 and IL8 at 6 hours, and IL1β and IL8 further increased by 24 hours, indicating a stronger pro-inflammatory response than the wild-type.

As differences between the wild-type and EPS-mutant strains were more pronounced after 24 hours, we conducted a broader analysis of their effects on vaginal cells inflammation using the Olink® Target 96 assay, which measures the production levels (fold change) of 92 inflammatory biomarkers (supplementary dataset). 49 of the 92 biomarkers tested were detected in most *in vitro* samples of vaginal epithelial cells (>55%) (**Fig. 4C**) and confirmed to be also present in human vaginal fluid when analyzing their prevalence in vaginal swabs of healthy female donors (**Fig. 4D**). Of these biomarkers, the presence of *L. crispatus* AMBV-0815 wild-type significantly enhanced the production of 11 biomarkers compared to non-treated vaginal epithelial cells: the cytokines IL6, IL8, the regulatory cytokines LAP TGF-beta-1, LIF; the chemokines CXCL1, CXCL10, CXCL11, CCL3 and CCL20; the growth factor FGF19 and tumor suppressive molecule CST5 (Cystatin D). In contrast, when the EPS layer was absent, as in the EPS-mutant, the cytokine CSF-1, the growth factor GDNF and the matrix metalloproteinase MMP-10 were upregulated, while the induction of LAP TGF-beta-1 and CST5 observed in the wild-type was no longer present, indicating a role of EPS in modulating these vaginal epithelial cells biomarkers (**Fig. 4C**).

### The presence of EPS modulates *L. crispatus* interactions with a 3D vaginal model

To better resemble the *in vivo* physiological conditions, a three-dimensional (3D) vaginal biomimetic model (adapted from Edwards et al 2022 ^32^) was used to investigate the role of cell wall EPS in *L. crispatus* host-interactions. The basolateral compartment was coated with collagen and supported a layer of fibroblast cells, resembling the connective tissue of the vaginal mucosa, which remains in contact with nutrients. The apical compartment consisted of stratified immortalized vaginal epithelial cells cultured without media to simulate the air-exposed epithelial layer of the vaginal lumen, promoting cell stratification and differentiation. The correct multilayered structure of the 3D model was assessed using light microscopy (**Supplementary Information, Fig. S7**). Subsequently, the model was exposed to *L. crispatus* AMBV-0815 wild-type and EPS-mutant strains, and the bacterial adhesion was monitored via two different imaging techniques, i.e. confocal microscopy and scanning electron microscopy (**Fig. 5**). Confocal microscopy enabled the visualization of a well-differentiated and morphologically intact culture (**Fig. 5a, b, c**), while scanning electron microscopy provided higher-magnification insights of the host surfaces and the adherent bacteria (**Fig. 5d, e, f**). As shown in **Fig. 5**, the EPS-mutant strain exhibited higher adhesion to the apical surface of the 3D vaginal model compared to the wild-type counterpart, consistent with observations from the vaginal cell monolayer.

**Figure 5.**
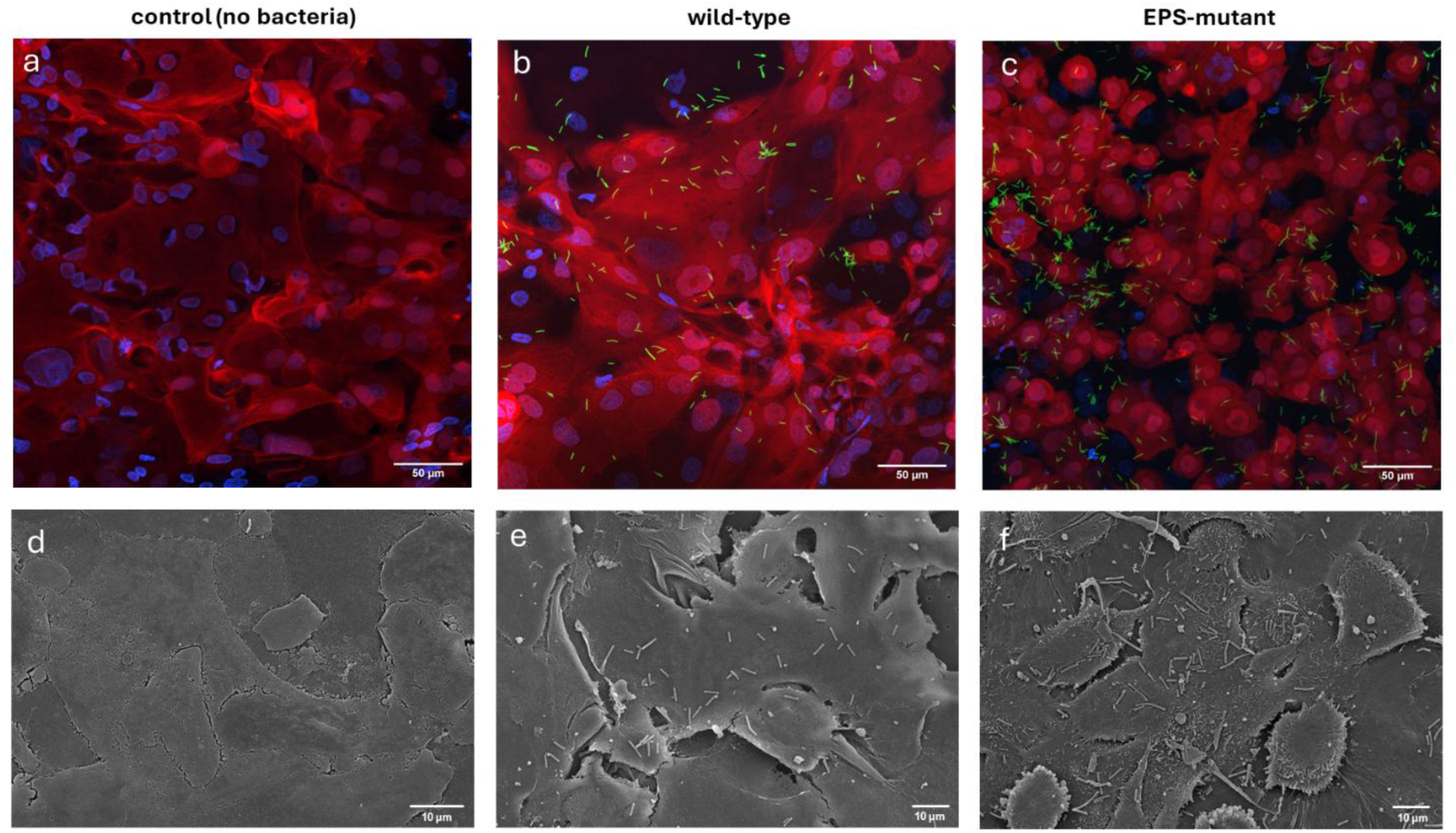
Confocal and scanning electron microscopy images showing the role of EPS in *L. crispatus* interaction with a 3D vaginal biomimetic model. For confocal microscopy (a,b,c), human cells were stained with Hoechst (nuclei, blue) and Deep Cell Mask (membranes, red), while *L. crispatus* AMBV-0815 wild-type and EPS-mutant strains were FITC-stained prior to adhesion. Images show the maximum intensity projection of (a) apical layer of differentiated vaginal cells without bacteria, (b) apical layer of vaginal cells with AMBV-0815 wild-type, and (c) apical layer of vaginal cells with EPS-mutant. For scanning electron microscopy (d,e,f), surfaces were gold-sputtered, and images depict (d) the apical layer of differentiated vaginal cells without bacteria, (e) the apical layer of vaginal cells with AMBV-0815 wild-type, and (f) the apical layer of vaginal cells with EPS-mutant. Scale bars are indicated.

To better understand the role of cell wall EPS in *L. crispatus* interactions with the vaginal immune system within a complex environment, we assessed the immunomodulatory effects of the AMBV-0815 wild-type and EPS-mutant strains using the Olink® Target 96 assay on both the apical and basolateral compartments of the 3D vaginal biomimetic model. Overall, the apical and basolateral sides exhibited distinct biomarker expression profiles (supplementary dataset). On the basolateral side, 45 of 92 biomarkers were detected above the LOD (>55%),but appeared unaffected by bacterial presence and were therefore not further analyzed. In contrast, on the apical side, directly exposed to bacteria, 56 of 92 biomarkers were detected above the LOD (>55%), with some showing different expression profiles depending on the bacterial strain. Notably, nearly all biomarkers identified *in vitro* were also prevalent in vaginal swabs from healthy female donors, confirming their physiological relevance (**Fig. 6A**). Interestingly, exposure to *L. crispatus* AMBV-0815 wild-type led to a mild increase in the production of the interleukin IL-17C and the chemokines CXCL5, CXCL6, and CCL20 compared to untreated vaginal epithelial cells. In contrast, the EPS-mutant did not alter the expression of CXCL5 and CXCL6 but did downregulate the costimulatory protein CD40 compared to the effect exerted by the wild-type (**Fig. 6B**). These results are consistent with differential immune interactions elicited by EPS-expressing versus EPS-deficient *L. crispatus* strain, even within a complex cellular milieu extending beyond vaginal epithelial cells.

**Figure 6.**
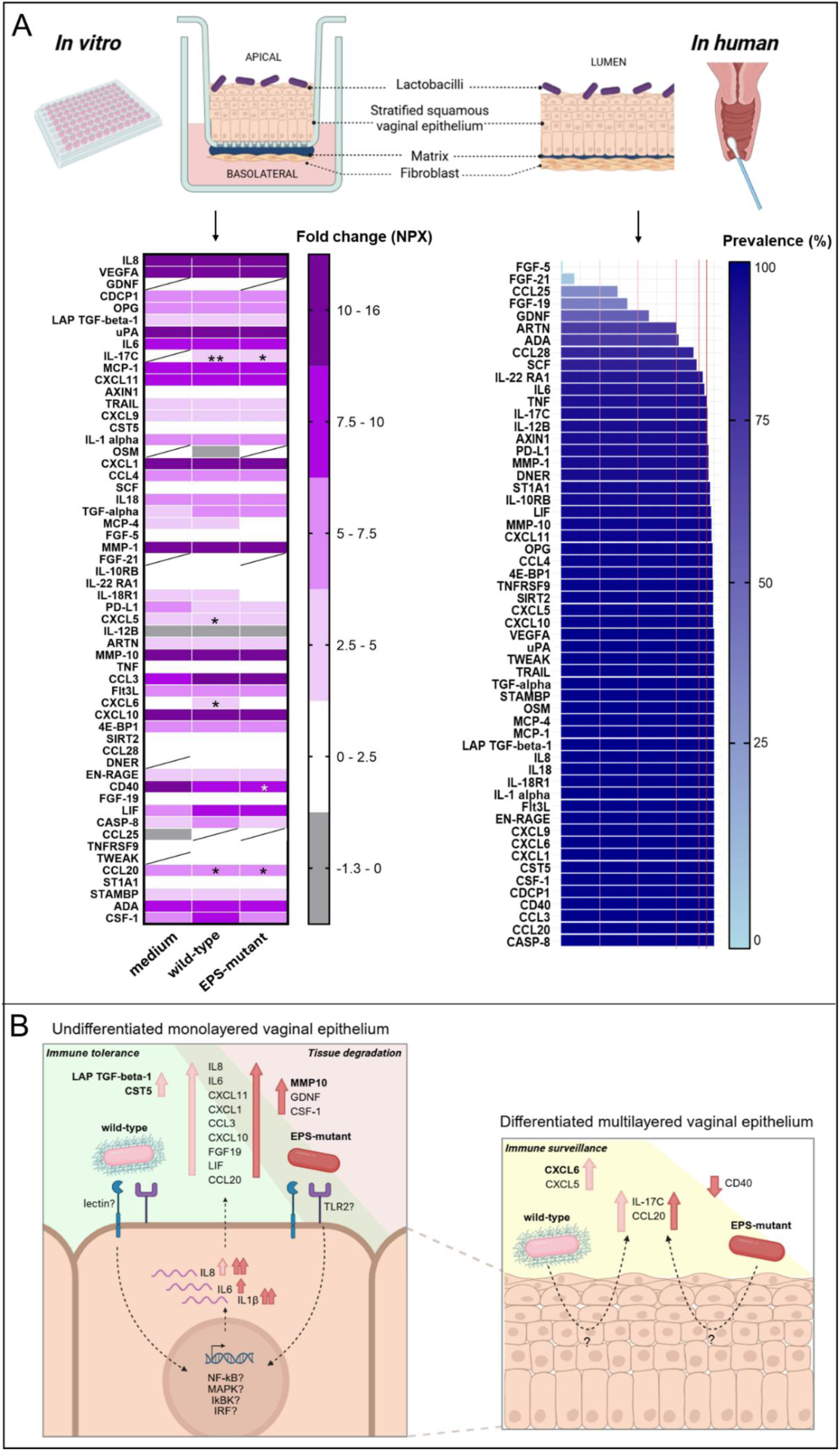
Validation of the EPS function in *L. crispatus* interactions with the vaginal immune system. (A) Effect of L. crispatus AMBV-0815 wild-type and EPS-mutant (24 h) on the inflammation panel of the 3D vaginal biomimetic model apical compartment (Olink® assay). KSFM cells medium was used as negative control. Data are reported as fold-change (NPX) of 56 inflammatory biomarkers. Biomarkers with more than 55% of samples below the limit of detection (LOD) represented by cells with a line. Statistical comparisons relative to the negative control are presented in the heat map (n=3). ANOVA two-way with Tukey correction (*p = 0.033, **p = 0.002, ***p < 0.001). Prevalence (%) of 56 immune biomarkers detected above LOD in human vaginal samples of healthy volunteers, as identified by the Olink® assay in multilayered immortalized vaginal epithelium. The horizontal lines indicate a 95%, 90%, 75%, 50% and 25% prevalence. (B) Schematic illustrating the role of cell wall *L. crispatus* EPS in modulating the immune response of immortalized vaginal epithelial cells (monolayer and multilayered in vitro models). Upward arrows indicate upregulation; downward arrows denote downregulation. NF-κB: Nuclear Factor kappa-light-chain-enhancer of activated B cells; MAPK: Mitogen-Activated Protein Kinases; IKBK: IκB Kinase Beta; IRF: Interferon Regulatory Factor. Unknown mechanism indicated with a question mark. Figure made with *Biorender.com*

## Discussion

The symbiotic relationship between vaginal *L. crispatus* and its host is a remarkable example of mutualism, where both organisms benefit and coevolve within the ecosystem. Exopolysaccharides (EPS) on the cell wall of lactobacilli modulate symbiosis and promote host tolerance in the gut, depending on the monosaccharide composition, structure, and thickness ^14^. As demonstrated in studies on genetic manipulable gut-adapted lactobacilli ^17,21,22^, the most observed phenotype is that EPS can mask (shield) bacterial molecules from host recognition, affecting their adhesion and immunogenicity on different cell lines. However, whether these effects apply to specific EPS molecules of *L. crispatus*, a dominant species in healthy women that is recalcitrant to cultivation and genetic modifications, remained an unanswered question. In this study, by integrating different approaches, including comparative genomics, citizen science datasets, mutagenesis, biochemistry, imaging, immunology, various vaginal models and validation with human samples, we were able to develop an innovative framework for the genetic and phenotypic analysis of protective EPS functions in *L. crispatus* on the vaginal mucosa.

We first identified an EPS biosynthetic gene cluster in a selection of vaginal *L. crispatus* strains available in-house, displaying a ‘generic’ organization in the genome. The first five genes (*epsA, epsC, epsD, epsB,* and *epsE*) were more conserved across strains than the remaining genes in the cluster. Pangenome analysis of the genus *Lactobacillus* revealed that these genes are highly conserved in *L. crispatus,* moderately conserved in *L. gasseri*, and only rarely present in *L. jensenii*. In *L. iners*, only *epsA* was found, indicating that this species is likely unable to produce this type of EPS. Of note, *epsE* was more variable, clustering into different orthogroups. Within this region, the priming glycosyltransferase *epsE*, widely recognized as the key initiator of EPS biosynthesis and encoding undecaprenyl-phosphate galactose phosphotransferase, exhibited lower sequence conservation than *epsA, epsC, epsD,* and *epsB*, which are implicated in EPS chain regulation ^16^. Moreover, *epsE* was less frequently detected as expressed in human samples, suggesting it may be subject to tighter regulatory control, potentially serving as a molecular switch for EPS production. In contrast, the downstream region of the cluster displayed greater sequence variability, particularly among genes encoding glycosyltransferases. This diversity likely underpins species- and strain-specific differences in EPS structure, including variation in monomer composition and branching patterns, and may contribute to functional specialization within the vaginal niche, although the lack of strict monosaccharide specificity in glycosyltransferases may limit how much these genetic differences affect EPS structure. Moreover, structural analysis of the capsular EPS from *L. crispatus* AMBV-0815 revealed multiple monomers, including ribose, a rare sugar among *Lactobacillaceae* ^33^. Most monosaccharides were also identified in previously characterized EPS from *L. crispatus* BC1 and BC5 ^34^, supporting a genetic–structural consistency across strains to some extent, despite differences in their relative abundance. These findings underscore the limitations of predicting EPS structure from gene content alone and highlight the need for broader structure characterization across vaginal *L. crispatus* strains.

To functionally analyze the role of EPS in host interactions without altering its native conformation presented on live cells, we created an isogenic *epsE* mutant in our competent model *L. crispatus* AMBV-0815 strain. We demonstrated that deleting the *epsE* gene led to a substantially decreased yield of capsular sugars, with lower levels of D-glucosamine and D-galactose, confirming a key role of this gene in producing capsular EPS enriched in these two monomers. A similar role for the priming glycosyltransferase was observed for EPS production in gut-derived lactobacilli such as *L. rhamnosus* GG ^17^, *L. plantarum* WCFS1 ^21^, *L. gasseri* DSM 14869 ^30^ and *Lacticaseibacillus paracasei* subsp. *paracasei* BGSJ2-8 ^35^. However, some capsular sugars were still detected in our *L. crispatus* EPS-mutant, suggesting that other glycosyltransferases may contribute to alternative pathways independent of the priming glycosyltransferase, such as those involved in teichoic acid, glycosylated protein, or peptidoglycan synthesis (for an overview of these glycoconjugates, we refer to ^14^).

We then demonstrated that cell wall EPS in *L. crispatus* AMBV-0815 supports its biofilm formation on abiotic surfaces by promoting bacterial cohesion, while reducing the adhesive capacities on both simple and complex vaginal cell models (VK2/E6E7 monolayer and 3D vaginal biomimetic model). This phenomenon could be explained by the reduced exposure of adhesive molecules mediated by EPS, which aligns with the shielding effect hypothesis previously proposed for gut lactobacilli EPS such as *L. rhamnosus* GG ^17^, *L. plantarum* Lp90 ^22^ and *L. johnsonii* FI9785 ^36^. In contrast, other studies on *L. paracasei* subsp. *paracasei* BGSJ2-8 ^35^, *L. plantarum* WCFS1 ^22^ and vaginal *L. gasseri* DSM14869 ^30^ showed that EPS can also be used by bacteria to adhere on epithelial cells, highlighting the diverse modulatory roles of these polymers in lactobacilli interactions depending on the producing species/strain, specific EPS structure and target adhesion surface tested.

Given that the EPS layer is not only itself a first contact point with target cells but also alters the exposure of underlying *L. crispatus* cell wall elements, we then explored whether reducing the EPS affects the presentation of immunogenic components and influences the human immune system. To investigate this, we used different cell lines, starting with the innate immune activation in human monocytes (THP1-Dual™) and assessing the immune response in vaginal cell models. *L. crispatus* AMBV-0815 wild-type triggered both NF-κB and IRF pathways in monocytes, which are transcription factors involved in cytokine production and type I interferon responses, crucial for antimicrobial defenses. These pathways have been shown before to be induced by commensal gut- and saliva-derived lactobacilli *L. rhamnosus* GG and *L. plantarum* WCFS1 respectively and linked to a host immune tolerance-effect ^37^. Notably, reducing the EPS layer in vaginal *L. crispatus* AMBV-0815 led to increased IRF pathway expression, suggesting that EPS prevents excessive immune activation, promoting tolerance and symbiosis. Such EPS-mediated immune protection aligns with findings in gut model probiotics *L. plantarum* WCFS1 ^21^ and SF2A35B ^22^, where HEK-293 immune cells showed higher NF-κB production in response to the EPS-mutants than wild-type strains. However, no such differences were observed for *L. plantarum* Lp90 ^22^, highlighting strain-specific EPS effects on immune and human host cells.

By implementing a high-throughput immune profiling technology (Olink®), we also investigated the role of EPS in the immune response of vaginal cells to *L. crispatus*, using both a monolayer and a 3D vaginal biomimetic model optimized for studying molecular microbe–host interactions. Our findings demonstrated that the immune biomarkers expression by vaginal cells is influenced by its cellular differentiation. Differentiated vaginal cells in the 3D vaginal biomimetic model exhibited increased production of chemokines (CXCL6, CXCL9, CCL28, CCL25) and the immune checkpoint molecule PD-L1 (Programmed Death-Ligand 1) compared to monolayers, emphasizing the role of cellular maturation in shaping immune signaling. Additionally, we demonstrated that EPS presence in the *L. crispatus* cell wall helps balance vaginal immune responses by modulating diverse immune pathways, with effects varying according to the cell model employed. On the vaginal monolayered epithelium, the presence of EPS on *L. crispatus* supported vaginal immune tolerance to these bacteria, balancing mucosal resilience and protecting against tissue degradation. The EPS layer on bacterial cells appeared to either directly or indirectly enhance the expression of LAP TGF-beta-1, the latent form of TGF-beta-1 (transforming growth factor beta), a cytokine essential for immunosuppression and tissue homeostasis ^38^. In the gut niche, TGF-beta-1 induction by the commensal bacteria *Clostridium* is a known mechanism associated with immune tolerance and maintenance of the mucosal equilibrium ^39^. Here, we present – to the best of our knowledge - the first report that vaginal lactobacilli, specifically through EPS, could also regulate the expression of LAP TGF-beta-1, proposing a previously underrecognized immunomodulatory function in the vagina. Furthermore, EPS-expressing *L. crispatus* was able to induce CST5, a cysteine protease inhibitor typically found in saliva and in the gut, critical for maintaining epithelial barrier integrity and antimicrobial defense ^40^. These two biomarkers warrant further investigation, as they are closely associated with key immunoregulatory functions, including the clearance of human papillomavirus (HPV) and the prevention of cervical cancer progression ^41,42^. On the other hand, the reduction of EPS in the mutant was associated with an over-expression of pro-inflammatory cytokines including IL1β, IL6, and IL8 in the vaginal monolayer epithelium, alongside the upregulation of MMP10, a matrix metalloproteinase involved in extracellular matrix degradation associated to bacterial vaginosis infections ^43^. In differentiated vaginal cells, EPS on *L. crispatus* seemed to support immune surveillance by mildly upregulating the chemokines CXCL5 and CXCL6, which bind to CXCR2 on immune cells, regulating B cell lymphopoiesis and neutrophil trafficking ^44^. In the absence of EPS, the *L. crispatus* mutant reduced CD40 expression, a key co-receptor involved in immune cell activation, potentially weakening the host’s defense against infections. Previous *in vitro* research has shown that vaginal *L. crispatus* upregulates CD40 and the differentiation of monocytic precursors into Langerhans-like cells ^45^. Our findings suggest that EPS may also play a critical and previously underappreciated role in host immune surveillance, considered a protective mechanism against vaginal infections ^28^. Furthermore, by applying Olink® technology to human samples, we validated that all immune biomarkers affected by the presence or absence of EPS were also significantly expressed in the vaginal environment of women.

Our findings show that the specific heteropolymeric EPS, enriched in D-glucosamine and D-galactose, from a vaginal *L. crispatus* strain actively modulates complex immune pathways when expressed on live cells, functioning as part of a ’whole-cell complex’ that orchestrates a balanced immune response. These results contribute functional insights to previous epidemiological studies on vaginal *L. jensenii* ^46^ and *L. crispatus* ^47^, which have shown through genome analyses that the complexity of surface glycans modulates host interactions, influencing immune recognition and bacterial persistence in the vaginal niche. Specific glycan branches and conformations within EPS can be recognized by human lectin receptors with different sugar specificity, impacting the host immune response in multiple ways ^48^. For example, glycans like mannose and glucose exposed on the S-layer proteins of *L. crispatus* have been shown to trigger an anti-inflammatory response via the lectin receptor DC-SIGN, contributing to S-layer-mediated shielding of TLR ligands ^25^. Although the glycosidic linkages and branching patterns in our strain remain uncharacterized, we hypothesize that immune modulation by *L. crispatus* EPS involves both direct lectin recognition and indirect shielding of underlying more pro-inflammatory molecules such as lipotechoic acid or lipoproteins, with the precise role of specific sugars still to be elucidated. Additionally, immune shifts observed in the absence of EPS across different cell models highlight the need for further investigation into its role within the dynamic mucosal environment. Furthermore, our finding that *L. crispatus* EPS genes and key host immune mediators identified *in vitro* are also expressed in the vaginal niche - detected in vaginal swabs from healthy female donors - highlights the physiological relevance of our results. We also acknowledge the limitations of our *in vitro* models and the complexity of host–microbe interactions *in vivo* and highlight that further mechanistic and clinical research is needed to dissect the specific molecular pathways involved, validate causality, and assess the broader relevance of EPS-mediated immune modulation across diverse populations and clinical contexts. Despite the valuable insights gained, our study was constrained by the limited genetic accessibility of vaginal *L. crispatus* isolates, which restricted genetic manipulation to a single strain. Expanding genetic tools and enabling optimization across multiple strains would be essential to more comprehensively evaluate the role of EPS at the species level. Furthermore, our findings emphasize the limitations of inferring EPS structure solely from gene content, underscoring the need for broader structural characterization across diverse *L. crispatus* strains. A more integrated approach, combining genomic, structural, and functional analyses, will be critical for elucidating strain-specific EPS features.

In conclusion, our integrative approach reveals that D-glucosamine– and D-galactose–enriched EPS from *L. crispatus* shapes host–microbe interactions in the vaginal mucosa by supporting biofilm architecture, limiting adhesion, and modulating immune responses. These findings highlight the presence of *L. crispatus* EPS as a key contributor to immune balance and epithelial homeostasis in the vaginal niche.

## Methods

### Strains and growth conditions

Lactobacilli strains used in the present study are listed in **Tab. 1**. For experimental analyses, the *L. crispatus* strains were grown in a microaerobic atmosphere, containing 5% CO_2_*. Escherichia coli* DH5α was grown in Lysogeny Broth (Difco) medium with aeration at 37°C. Media were supplemented with antibiotics (Sigma-Aldrich) when appropriate: chloramphenicol (Cm) 10 μg/mL or 25 µg/mL for lactobacilli or *E. coli*, respectively; Ampicillin (Amp) 100 μg/mL for *E. coli*. When preparing electrocompetent cells, MRS was supplemented with glycine 2% (w/v) (Fisher Scientific) and sucrose 2% (w/v) (Merck).

**Table 1.** Lactobacilli strains used in this study.

| Species | Strains | Origin | References | Annotation |
| --- | --- | --- | --- | --- |
| ***Lactobacillus***<br>***crispatus*** | BC1 | Human<br>vagina | 4 | Genome (NCBI)<br>GCA_040109495.1 |
|  | BC3 |  |  | Genome (NCBI)<br>ASM4134485v1 |
|  | BC5 |  |  | Genome (NCBI)<br>GCF_014654865.1 |
|  | AMBV-0006 | Human vagina | <sup>6</sup> | EPS gene cluster – this work (supplementary dataset) |
|  | AMBV-0815 |  |  |  |
|  | AMBV-1513 |  |  |  |
|  | AMBV-2491 |  |  |  |
| | AMBV-0815Δ*epsE* | AMBV-0815 *epsE* knockout mutant; *epsE::cat(M)* | This work | - |
| *Lactiplantibacillus plantarum* | WCFS1 | Human saliva | <sup>49</sup> | Genome (NCBI) GCF_000203855.3 |

### Comparative genomic and pangenome analysis

The genome of 7 in-house *L. crispatus* strains (see **Tab. 1**) was analyzed for the presence of the priming glycosyltransferase gene (*epsE*), involved in the main step of the EPS biosynthesis in lactobacilli. To identify the *epsE* gene in *L. crispatus* strains, the aminoacidic sequence of vaginal *L. gasseri* DSM14869 priming glycosyltransferare gene (Accession: N506_0400 gene) ^30^ was used as a query for a tblastn against the genome *Lactobacillus crispatus* BC1 strain and hit > 50% identity was considered as valid. Afterwards, *L. crispatus* BC1 *epsE* was used as reference to identify the *epsE* in other selected *L. crispatus* strains and hits > 85% identity were considered as valid. The EPS gene cluster of the 7 in-house *L. crispatus* strains was investigated using the CLC Genomics Workbench Version 24.0.1 (Qiagen, Hilden, Germany) and sequence homologies between the gene clusters were identified by alignments.

All publicly available *Lactobacillaceae* genomes (including *L. crispatus* BC1, BC3 and BC5 genomes, see **Tab. 1**) were downloaded from the Genome Taxonomy Database (GTDB, gtdb.ecogenomic.org, version r220 ^50^) and supplemented with in-house genome-sequenced strains (*L. crispatus* AMBV-0006, AMBV-0815, AMBV-1513, AMBV-2491). With checkM, any incomplete (<95%) or contaminated (>5%) genomes were excluded. A core genome tree was inferred for all representative *Lactobacillaceae* genomes with SCARAP as described in Eilers et al. ^51^ on 296 core genes. The tree was subsequently filtered ^52^ on the genus *Lactobacillus*. This resulted in a dataset of 2317 public genomes. To avoid pseudoreplication, strains with an average nucleotide identity (ANI) cutoff of 99.99% were clustered together, retaining 1609 genomes with ANI values below this cutoff. All genes were predicted with Prodigal ^53^ and the pan genomes of this family were inferred using the SCARAP tool ^54^. For visualization purposes, a core gene tree was constructed with orthogroups of detected genes in *L. rhamnosus* GG, *L. plantarum* WCFS1, *L. crispatus* BC1, *L. crispatus* BC5, *L. crispatus* AMBV-0815, *L. gasseri* DSM14869, *L. johnsonii* FI9758 and *L. reuteri* IRT: LytR transcriptional regulator (*epsA*), tyrosine kinase modulator (*epsC*), tyrosine kinase (*epsD*), phosphotyrosine phosphatase (*epsB*) and the priming glycosyltransferase (*epsE*) (supplementary dataset). Tree and heatmap was visualized in R with ggtree and tidygenomes packages.

### Transcriptomic analysis

Raw metatranscriptomes of the 144 samples from the study with accession number PRJEB50547 were downloaded from ENA ^31^. Adapter trimming and quality filtering of the reads was performed using fastp (version 0.20.1) with default settings ^55^. Afterwards, a first pass of human read removal was performed with the hostile tool (version 2.0.0) using the default index ^56^. Then an index was created using the EPS genes of interest (*epsA, epsD, epsC, epsB, epsE*) of *L. crispatus* strains (see **Tab.1**), the VIRGO vaginal non-redundant gene catalog ^57^ and the human transcriptome index from hostile to map the reads with version 1.10.3 of the salmon tool ^58^. The resulting individual transcripts per million (TPM) counts were merged in a count table and visualized using the pandas and seaborn python packages. The *Lactobacillus crispatus* 16S rRNA transcript was used as a marker for *L. crispatus* presence and the 19 samples not containing any of the markers were discarded.

### Mutagenesis approach in *L. crispatus*

A double homologous recombination strategy was used in this study for deleting *epsE* in *L. crispatus.* Plasmids, primers, and strains for genetic manipulation used are listed in **Tab. 2**. Gene deletion was achieved by chromosomal replacement of *epsE* with the antibiotic selection marker gene *cat* (encoding chloramphenicol acetyltransferase), following the strategy reported in Lebeer et al., 2009 ^17^, with minor modifications. The *L. crispatus* BC1 genome was used for primer design to amplify the up- and downstream regions of *epsE*. Q5 High-Fidelity DNA Polymerase (New Englands Biolabs) was used to obtain fragments of DNA to be cloned into the suicide vector pUC19. DNA segments of 528 bp and 567 bp up- and downstream of *epsE* were amplified by PCR using primer pairs with overhang sequences HU-F(PstI)/HU-R and HD-F/HD-R(SacI), respectively. Similarly, *cat* under the control of the constitutive promoter p32 was obtained from plasmid pNZ5319 by amplification with primer pair Cat-F/Cat-R. Subsequently, plasmid pUC19 was digested with restriction enzymes PstI and SacI (Anza™, ThermoFisher). The PCR amplicons and the linearized plasmid were purified using the NucleoSpin Gel and PCR Clean-up kit (Macherey-Nagel). After purification, a Gibson Assembly (New England Biolabs) was performed to ligate the three fragments into the linearized plasmid pUC19. The ligation mixture was then transformed into *E. coli* DH5α, followed by plating with Amp and Cm as selection markers. The resulting plasmid pAMB5815 was purified from a colony using the Monarch Plasmid Purification Maxiprep Kit (New England Biolabs) after sequence confirmation by Sanger sequencing using primer pairs M13-F/HD_left-R and HU_right-F/M13-R.

**Table 2.**
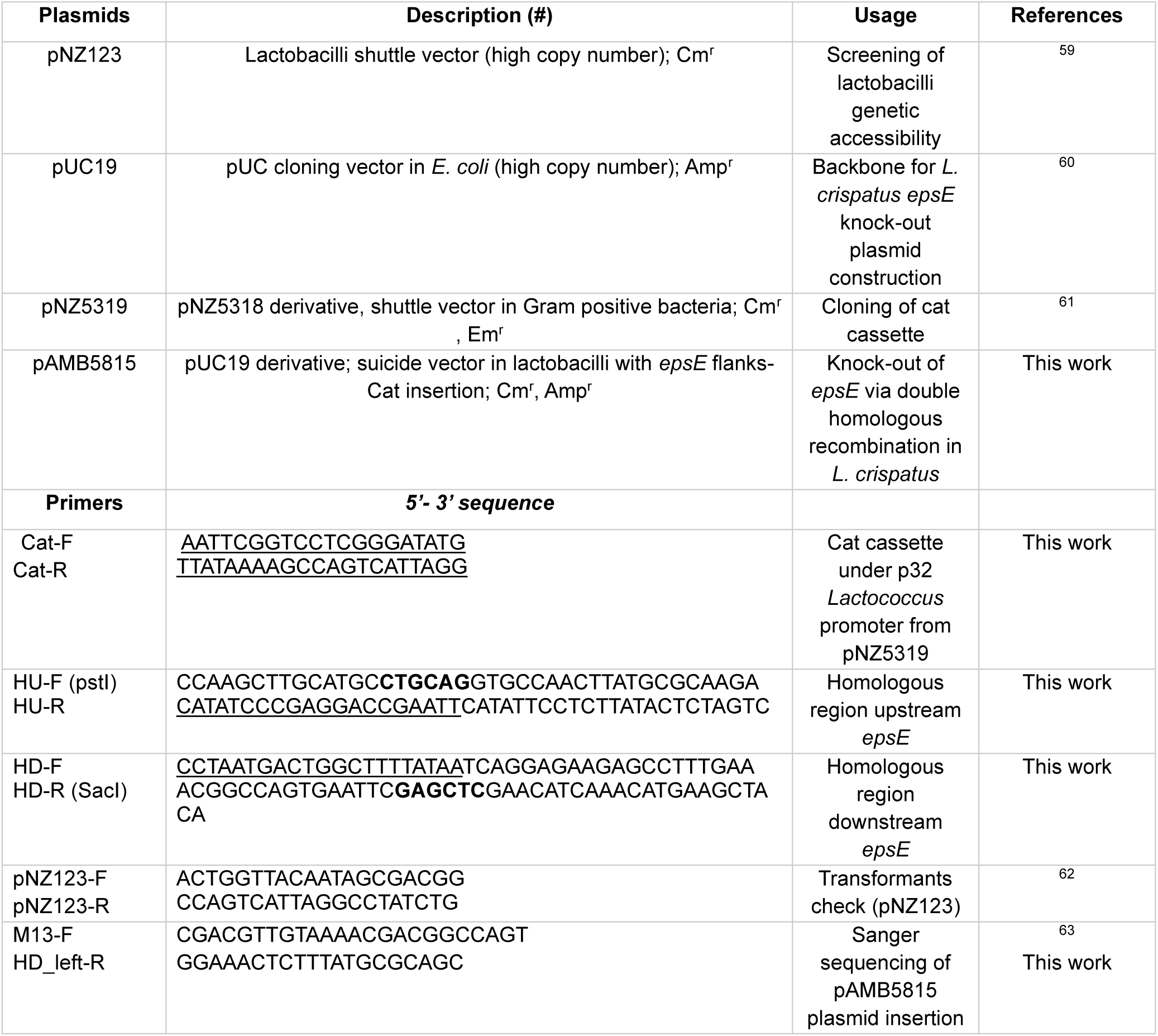
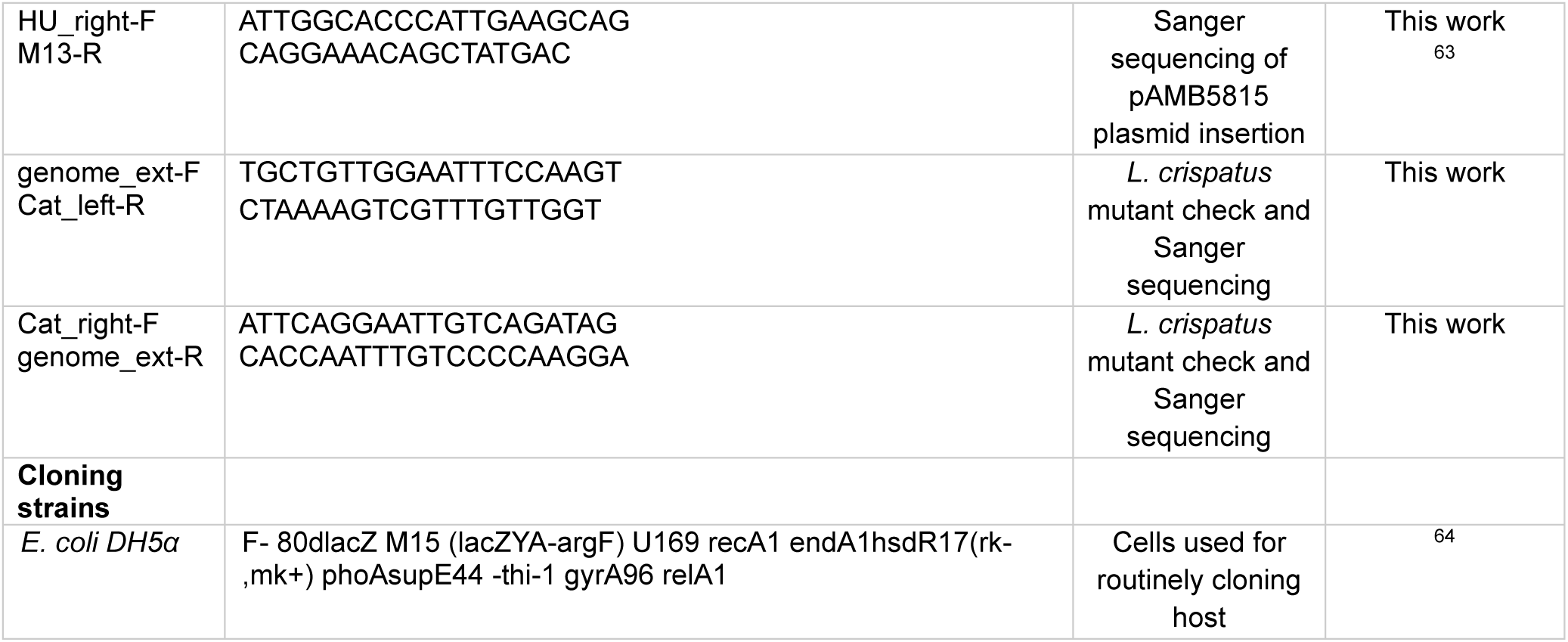
List of plasmids, primers and strains used for genetic manipulations in lactobacilli. Complementary sequences between primers are underlined. Restriction sites in the primers are indicated in brackets and sequences marked in bold. (#) Cm^r^, chloramphenicol resistant; Amp^r^, ampicillin resistant; Em^r^, erythromycin resistant.

### Transformation efficiencies and *epsE* mutant identification in *L. crispatus* strains

The electrocompetent cells of *L. crispatus* strains listed in **Tab. 1** were prepared by using the Mason et al., 2005 protocol, with some modifications. As a model strain for genetic manipulation *Lactiplantibacillus plantarum* WCFS1 was used due to its high competency with glycine-sucrose growth-based protocols. In detail, an overnight culture of an *L. crispatus* strain was diluted 1:100 in 10 mL of fresh MRS supplemented with glycine 2% (w/v) and sucrose 2% (w/v). After 18h of incubation, 1x10^4^; 1x10^5^ and 1x10^6^ dilutions were prepared in 50 mL of fresh MRS supplemented with glycine 2% (w/v) and sucrose 2% (w/v). After ca. 16h incubation, OD_600_ was measured and the dilution in the range of OD600 0.4-0.6, corresponding to 1.6-2.4x10^9^ CFU/mL, was selected for the next steps which were performed on ice. The cells were harvested by centrifugation (4000x g, 10 min, 4°C) and competent cells of lactobacilli were prepared by washing the cells twice with cold distilled water and finally resuspended in 5 mL of cold 50 mM EDTA (Sigma-Aldrich) for 5 minutes. After incubation, a volume of 20 mL of cold distilled water was added and the cells were harvested by centrifugation as mentioned before. The pellet was washed once with 2 mL of cold sucrose 300 mM. The competent cells were harvested by centrifugation (1370x g, 10 min, 4°C) and resuspended in 1 mL of cold sucrose 300 mM. Cells were aliquoted (200 μL) and transformed with 1-2 µg of plasmids pNZ123 or pAMB5815, or 2 μL of ultrapure water (negative control) using the Gene Pulser Electroporator (Bio-Rad, Hemel Hempstead, UK) in 0.2-cm electrocuvette Gap (Bio-Rad, Hemel Hempstead, UK) with the following pulse settings: an exponentially decaying pulse of 1.5kV, 200Ω parallel resistance, and 25 µF capacitance. Electroporated cells were immediately diluted 1:5 in 1 mL with pre-warmed MRS media and incubated for 3h at 37°C, 5% CO_2_. After incubation, cells were plated on MRS agar (1.5% w/v) supplemented with Cm and incubated for 72h. Transformation efficiencies (TE) were calculated using the formula (1):

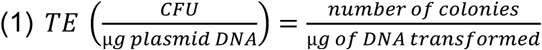

After 3 days of incubation on MRS plates containing Cm, colonies representing candidate transformants (pNZ123) were checked for the presence of the replicative plasmid using via PCR using the primer pairs: pNZ123-F/pNZ123-R. Colonies representing mutants (pAMB5815) were screened for the double homologous recombination event to have occurred by PCR and Sanger sequencing using the primer pairs: genome_ext-F/Cat_left-R, Cat_right-F/genome_ext-R and genome_ext-F/genome_ext-R (**Tab. 2, see Supplementary Information, Fig. S2**).

*L. crispatus* AMBV-0815 resulted to be knocked out of *epsE* and the mutant strain, named *L. crispatus* AMBV-0815Δ*epsE* (EPS-mutant) was used to perform all the phenotypic and functional assays reported below.

### Growth kinetics

To assess the effect of *epsE* deletion on *L. crispatus* growth kinetics, the growth of *L. crispatus* AMBV-0815 wild-type and EPS-mutant was monitored for 24h. Briefly, lactobacilli suspensions were prepared at 1 × 10^6^ CFU/mL in MRS medium. A volume of 200 μL was then inoculated into each well of a 96-multiwell plate and the absorbance (ABS-OD_600_) was read in continuous mode for 24h by using the Synergy HTX Plate Reader (BioTek). The cells growth rate (*μ*), which indicated the rate at which the cell population increases, was then calculated following the formula (2):

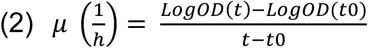

Where “t” indicates the final time point and “t0” indicates the initial time point of the exponential phase during bacterial growth. Each condition was repeated 3 times.

### Transmission electron microscopy analysis

To visualize the polysaccharides structure on the cell wall, *L. crispatus* AMBV-0815 wild-type and EPS-mutant cells were negatively stained with uranyl acetate and analyzed with a Tecnai G2 Spirit Bio TWIN microscope (Fei, Europe BV, Zaventem, Belgium) at 120 kV transmission electron microscopy (TEM), as previously reported ^17,66^. The quantification of the amount of surface polysaccharides on AMBV-0815 wild-type was conducted by analyzing at least 3 TEM micrographs with ImageJ f54, version Java 1.8.0 (National Institute of Health, NIH).

### Capsular sugars isolation

Capsular sugars were isolated from *L. crispatus* AMBV-0815 wild-type and EPS-mutant cells as previously described, with some modifications ^67^. Briefly, cells were harvested from 500 mL overnight cultures by centrifugation (8000x g, 10 min, 4°C), and washed twice with sterile saline (8000x g, 10 min, 4°C). The cell pellet was resuspended in 25 mL of 50 mM EDTA (pH 8) (Sigma-Aldrich), and the suspension was stirred gently (100 rpm) for 4h, at 4°C. After centrifugation (15294x g, 20 min, 4°C), two volumes of chilled ethanol (99%) (Merck) were added to the supernatant, and the mixture was incubated at 4°C overnight to permit EPS precipitation. After subsequent centrifugation (15294x g, 40 min, 4°C), the pellet (containing EPS) was resuspended in 10 mL of sterile distilled water and the suspension was dialyzed against 5 L of distilled water using a 6–8 kDa dialysis membrane (Spectra/Por, VWR International, Radnor, Pennsylvania, USA) for 2 days, with 3 water changes per day under agitation. Afterwards, proteins were removed from the EPS samples following the protocol reported in Allonsius et al., 2017. Briefly, proteins were precipitated in trichloroacetic acid 20% (w/v) (Sigma-Aldrich) for 2h under mild agitation (100 rpm) at 4°C, and removed by centrifugation (15294x g, 40 min, 4°C). The trichloroacetic acid was removed by dialysis against 5 L of distilled water for three days with three changes of water under agitation. The purified solution was freeze dried at - 47°C and 0.01 atm (Christ Freeze Dryer ALPHA 1–2, Milan, Italy). The amount of crude sugars from *L. crispatus* AMBV-0815 wild-type and EPS-mutant was weighted, and yields were normalized for 10^9^ CFU. Each condition was repeated 2 times.

### Monosaccharides composition

The monosaccharide composition of crude sugars was defined through Liquid Chromatography-Mass Spectrometry (LC-MS), following the procedure described below. **(i) Sample preparation**. Freeze-dried sugars (1 mg per sample) were hydrolyzed in 250 μL of HCl 4 M and incubated 1h at 99°C under gentle shaking (300 rpm, Thermomixer Comfort). The samples were then neutralized with 250 μL of NaOH 4 M. Subsequently, monosaccharides underwent derivatization by mixing 120 μL of the hydrolyzed solutions with 180 μL of 0.5 M NaOH. Then, 200 μL of this mixture was combined with 200 μL of PMP (0.5 M in methanol) and incubated at 70 °C for 1h with gentle shaking (300 rpm). After cooling to room temperature, 200 μL of HCl 0.3 M and 300 μL of Tris buffer (1.5 M, pH 7) were sequentially added for neutralization. The excess PMP was removed by extracting the mixtures three times with 500 μL of dichloromethane. The samples were aliquoted and stored at −20°C until analysis. Standard solutions (50 mM) of D-Glucosamine, D-Mannose, D-Galactosamine, D-Ribose, D-Rhamnose, D-Glucose, D-Galactose, D-Xylose, L-Fucose, N-Acetyl Glucosamine, D-Arabinose, D-Glucuronic Acid, N-Acetyl Galactosamine, Lactose, and D-Galacturonic Acid were derivatized using the same procedure. This entire process was performed in triplicate for each sample. **(ii) Quantitative analysis.** To build calibration curves, the 0.74 mM solution of each derivatized monosaccharide was diluted with sodium acetate buffer 100 mM pH 4 to get working solutions ranging from 0.00122 to 6.25 μM for D-Glucosamine, D-Mannose, D-Galactosamine, D-Ribose, D-Rhamnose and D-Glucose, from 0.00244 to 6.25 μM for D-Galactose and L-Fucose and from 0.0244 to 0.781 µM for D-Xylose. Standard solutions were analysed by LC–MS method reported below. The limit of detection (LOD) was determined through LC-MS analysis of serially diluted standard solutions and was defined as the concentration at which a signal-to-noise (S/N) ratio of 10 was reached. For the quantitation of monosaccharides, samples were diluted 1:10 with ammonium acetate buffer 100 mM, pH 4. **(iii) LC-MS analyses**. LC-MS was performed on a UPLC system (Acquity, quaternary Solvent Menager) coupled to a quadrupole–time-of-flight hybrid mass spectrometer (Xevo G2-XS QTof, Waters) equipped with an electrospray ionization interface operating in positive ion mode. Chromatographic separation was carried out on an Acquity UPLC BEH C18 column (130 Å, 1,7 μm, 2,1 mm x 150 mm) thermostated at 65°C. The analysis was conducted under isocratic conditions using a mobile phase consisting of 100 mM ammonium acetate buffer (pH 4) and acetonitrile (82:18, v/v). The flow rate was set at 0.6 mL/min, and the injection volume was 5 µL.

The mass spectrometer operated in high sensitivity mode using a capillary voltage of 0.8 kV and a cone voltage of 40 V. Cone and desolvation gas flow were 50 and 1200 L/h, respectively, while source and desolvation gas temperature were 150 and 600°C, respectively. Leucine enkephalin (0.1 ng/μL) was used as lock mass (m/z 556.2771). To identify the monosaccharide data were acquired in MS^E^ mode from *m/z* 50 to 1200, creating two discrete and independent interleaved acquisition functions. The first acquisition, set at 6.0 eV of collision energy, collected unfragmented data, while the second had a collision energy ramp from 20–30 eV and collected fragmented data. Argon was used for collision-induced dissociation. Unifi software (Waters, Manchester, UK) was used for visualization and alignment of low- and high-energy information.

For the quantitative analysis acquisition was performed in Total ion Current (TIC) mode in the *m/z* range 450-750. Peak area of standard PMP-monosaccharide was derived from the corresponding extract ion chromatogram and was plotted against the concentration to obtain the calibration curve.

### Quantification of biofilm formation

*L. crispatus* AMBV-0815 wild-type and EPS-mutant were first inoculated in 10 mL MRS medium and cultured as reported above. Bacterial suspension was then diluted 1:100 in 10 mL of fresh MRS medium and incubated again at 37°C overnight. Biofilm formation assay was assessed following the protocol reported in Giordani et al 2023, with some modifications ^34^. Briefly, the overnight culture was diluted to 1x10^7^ CFU/mL in fresh MRS medium. Then, 200 μL was added to each well of a 96 multi-well round-bottom plates (Corning Inc., Pisa, Italy). MRS medium was used as negative control. The plate was then incubated at 37°C for 72h to allow biofilm formation. After incubation, the bacterial suspension was removed and the biofilm formed at the bottom of the wells was gently washed once with phosphate buffer saline (PBS) and immediately stained using 0.1% (v/w) crystal violet (Merk, Milan, Italy) for 30 min at room temperature. After washing once with sterile water, the bound crystal violet was solubilized in 200 μL of acetic acid 30% (v/v) (Merk, Milan, Italy) and the absorbance was measured at 595 nm (EnSpire Multimode Plate Reader, PerkinElmer Inc., Waltham, MA, USA). Biofilm formation values of the EPS-mutant were normalized for *L. crispatus* AMBV-0815 wild-type. Each condition was repeated 3 times.

### Cell lines and culturing

The human vaginal VK2/E6E7 cell line (ATCC, CRL-2616) was maintained in Keratinocyte-serum free (KSFM-Gibco) medium supplemented with 0.1 ng/mL human recombinant epidermal growth factor (hrEGF- Gibco), 0.05 mg/mL bovine pituitary extract (BPE-Gibco) and 0.4 mM CaCl_2_ at 37°C in 5% CO_2._ Every 3 days, for maintenance, the VK2/E6E7 cells were split 1:5-1:7 with optional addition of 100 U/mL of Pen-strep to the complete medium.

The THP1-Dual™ human monocyte cell line (Invivogen, thpd-nfis) was maintained in Roswell Park Memorial Institute (RPMI) 1640 (Gibco) medium supplemented with 25 mM HEPES (ThermoFisher Scientific), 10% heat-inactivated Fetal Calf Serum (hi-FCS) (GE Healthcare) and 2 mM L-glutamine at 37°C in 5% CO_2_. The cells were maintained in RPMI1640 with L-glutamine, HEPES, 10% (v/v) hi-FCS, 50 mg/mL Normocin (Invivogen) and 100 U/mL Pen-strep (Gibco), and split without exceeding a concentration of 2×10^6^ cells/mL. To maintain selection pressure, 10mg/mL blasticidin (Invivogen) and 100 mg/mL zeocin (Invivogen) were added to the culture medium every other passage.

The human fibroblast BJ cell line (ATCC, CRL-2522) was maintained in Eagle’s Minimum Essential Medium (EMEM- ATCC) supplemented with 10% fetal bovine serum (FBS) (Gibco) at 37°C in 5% CO_2_. Every 3 days, the BJ cells were split 1:5-1:7 with fresh medium.

### Adhesion assays on vaginal monolayers and fluorescence imaging

Adhesion rates of *L. crispatus* AMBV-0815 wild-type and EPS-mutant to vaginal cells were performed by using two assays: direct count ^69^ and indirect plating out ^70^. Human vaginal VK2/E6E7 cells were seeded at 5×10^4^ cells/cm^2^ with KFSM medium without antibiotics in 24-wells cell culture plates (Sarstedt AG & Co., Nümbrecht, Germany) provided with a sterile round coverslip (direct count) or not (indirect plating out). After ca. 3 days, when cells reached 80-90% confluency, the exhaust medium was replaced with fresh complete medium (0.2 mL per well) and bacterial suspensions were added to the monolayer, applying a MOE (multiplicity of exposure) of 100:1 (bacteria: VK2/E6E7 cells). Plates were incubated for 3h, at 37°C in 5% CO_2_. For the direct count of bacteria adherent to VK2/E6E7 cell monolayers, samples were stained with May-Grunwald/Giemsa as reported before ^71^ and counted using a Nikon Eclipse 21 microscope (Objective 100×, Nikon, Tokyo, Japan) considering at least 20 microscopic fields (100 human cells). The ratio of bacteria/VK2 cells was calculated. For the indirect count, adherent bacteria were washed twice with PBS and detached with trypsin 0.5% (w/v) (Promocell). Afterwards, serial dilutions in PBS were plated out on MRS agar (1.5% w/v). Percentages of adherent bacteria compared to the initial inoculum were calculated. For both assays, the adhesion values of the EPS-mutant (ratio of adherent bacteria or percentages of adhesion) were standardized to the one obtained for *L. crispatus* AMBV-0815 wild-type.

For fluorescence imaging acquisition, bacteria were fluorescence-stained previous adhesion with fluorescein isothiocyanate isomer I (FITC – Merck) as reported in Spacova et al., 2024 ^72^. Briefly, an amount of 1x10^8^ CFU was resuspended in 200 μL of FITC solution (final concentration 0.1 mg/mL) and incubated for 1h at 37°C, shaking (250 rpm). After centrifugation, stained bacteria were washed once with PBS and added to the VK2/E6E7 cells monolayer (grown on coverslips), following the MOE as reported above. Afterwards, bacteria adherent to the monolayer were washed twice in PBS to remove non-adherent bacteria, and images were recorded with the Leica DMi8 fluorescence microscope by visualizing FITC-fluorescence of the stained bacterial cells coupled with phase contrast modality for the vaginal cells.

### Survival experiments

To exclude potential direct or indirect effects of vaginal epithelial cells on bacterial viability after 3 h of exposure, we performed a bacterial survival assay. Briefly, a monolayer of vaginal epithelial cells (VK2/E6E7) was cultured on the apical side of 24-well ThinCert™ inserts (CELLSTAR®, Greiner Bio-One). Specifically, 1.5 × 10⁵ cells were seeded per insert and cultured for approximately 7 days at 37°C and 5% CO₂, with 500 µL of fresh medium added to the basolateral compartment. After 7 days, monolayer integrity was confirmed by measuring transepithelial electrical resistance (TEER), which was required to be ≥400 mΩ·cm², as previously reported ^73^.

Subsequently, a bacterial suspension was prepared in cell culture medium (KFSM) at a final concentration of 2.5 × 10⁶ CFU/mL. To ensure a total inoculum of 1 × 10⁷ CFU, 200 µL of the suspension was added to both the apical and basolateral compartments. The plates were then incubated for 3 h at 37°C and 5% CO₂. Following incubation, bacterial cells were recovered from the apical compartment by trypsinization (0.25% trypsin, 5 min, 37°C, 5% CO₂) and from the basolateral medium. Serial dilutions of both the initial inoculum and recovered bacteria were plated on MRS agar and incubated for 24 h at 37°C and 5% CO_2_. Colony-forming units (CFU) were enumerated, and bacterial survival was calculated for each compartment using the following formula (3):

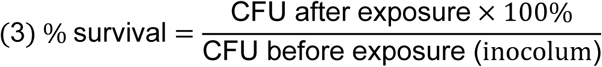

### Induction of NF-κB and IRF in THP1-Dual cells

The activation of NF-κB and IRF pathways in THP1-Dual™ reporter monocytes co-incubated with UV-inactivated bacteria was analyzed to investigate the immunomodulatory effect of the bacteria, as previously reported, with slight modifications ^37^.

Briefly, overnight bacterial cultures were washed once in PBS and resuspended at 2x10^7^ CFU/mL in RPMI 1640 medium with L-glutamine supplemented with 25 mM HEPES buffer and 10% (v/v) FBS without antibiotics. The bacterial suspensions were UV inactivated by 3 cycles of 15-min UV light irradiation followed by vortexing steps in a biosafety cabinet level 2. The UV inactivation was checked by spotting 5 μL of suspension on MRS agar plates, incubated overnight at 37°C in 5% CO_2_. THP1-Dual™ cells suspension was prepared with a concentration of 1x10^6^ cells/mL in RPMI 1640 medium (ThermoFisher Scientific) with L-glutamine supplemented with 25 mM HEPES buffer (Gibco) and 10% (v/v) FBS without antibiotics and seeded at 100 µL/well of a 96-well plate. Bacterial suspensions (2x10^7^ CFU, corresponding to a MOE of 20:1) were added per well and the plate was incubated for 20-24 hours at 37°C, in 5% CO_2_. The NF-κB activation was quantified by adding the para-nitrophenylphosphate (pNPP) buffer to the cell culture supernatants (ratio 2:1) and by analyzing the SEAP reporter activity at 405 nm with the Synergy HTX Plate Reader (BioTek). The activation of IRF was quantified by adding the QUANTI-Luc™ (InvivoGen) buffer to the supernatants (ratio 1:1) and by analyzing the luciferase reporter luminescence activity with the Synergy HTX Plate Reader (BioTek). As positive control, 10 ng/mL of lipopolysaccharides (LPS) from *E. coli* (Sigma) was used. Each condition was repeated 3 times.

### Immune response in vaginal monolayers

Human vaginal VK2/E6E7 cells were seeded in KFSM medium without antibiotics into a 12-well plate at a concentration of 4.2×10^4^ cells/cm^2^. After ca. 3 days, when cells reached 80-90% of confluency, the cells were exposed for 6h or 24h to *L. crispatus* AMBV-0815 wild-type or EPS-mutant suspended in VK2/E6E7 cells medium without antibiotics following the MOE reported above. As negative control, the cells were incubated with KFSM medium without antibiotics.

The expression of immune biomarkers by vaginal cells after 6h and 24h of bacterial exposure was assessed by reverse transcription (RT)-q-PCR. Briefly, the suspension was removed and the VK2/E6E7 cells were washed 3 times in PBS (Gibco). Afterwards, the total RNA was extracted with the RNeasy kit (Qiagen, Canada) and quantified using the Nanodrop spectrophotometer (Thermo Scientific). Afterwards, 500 ng of RNA were reverse transcribed into cDNA with SuperScript kit using the manufacture’s instruction (Invitrogen) and qPCRs were performed. In detail, qPCR was performed using StepOne Plus™ Real-Time PCR System (Thermo Fisher Scientific) with Power SYBR Green Master mix (Fisher Scientific) by using the primer pairs reported in **Tab. 3** and the cycle reported in **Tab. 4**. Data were normalized for the expression of two human housekeeping genes, *CYC1* and *GAPDH*. Each condition was repeated 3 times.

**Table 3.**
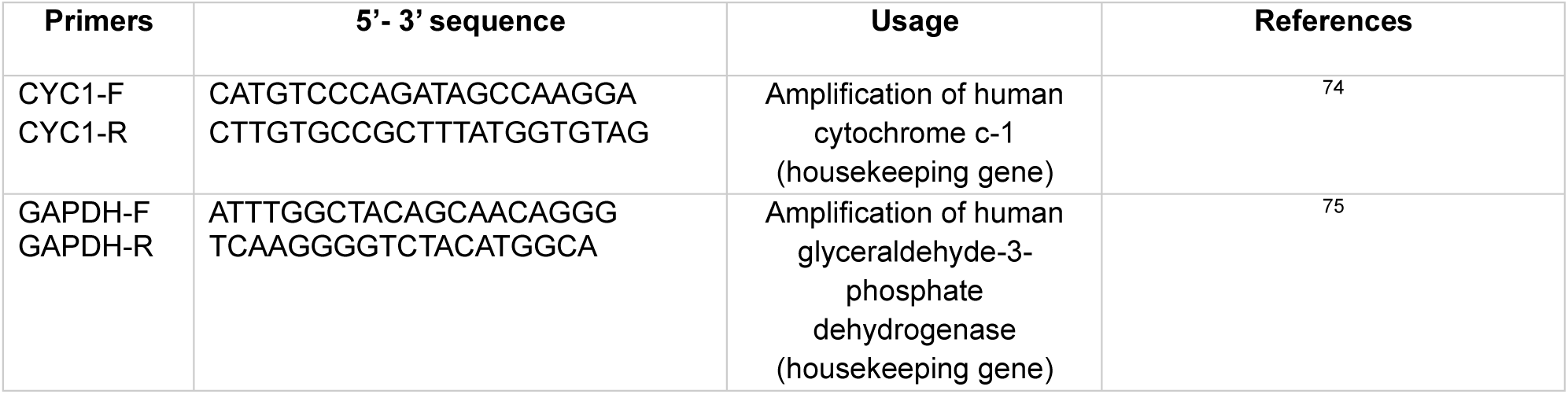
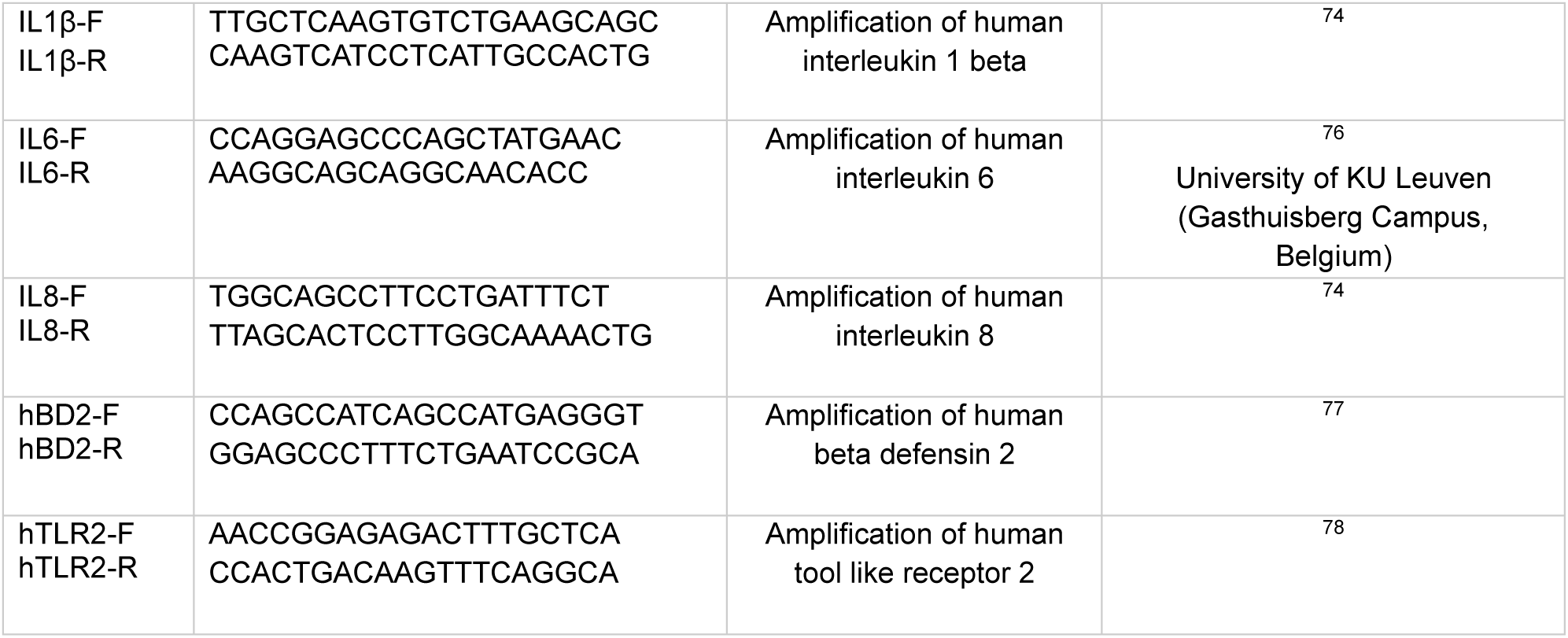
List of primers used for reverse transcription qPCR on VK2/E6E7 cell monolayer cDNA.

**Table 4.** Thermocycle for qPCR used in this study.

| qPCR stage | Temp. (°C) | Time |  |
| --- | --- | --- | --- |
| Pre-incubation | 50 | 10' |  |
|  | 95 | 20" |  |
| Denaturation | 95 | 3" | 40 cycles |
| Amplification | 60 | 30" |  |
| Melting curve | 95 | 15" |  |
|  | 60 | 1' |  |
|  | 95 | 15" |  |
| Cooling | 40 | 10" |  |

The inflammation panel of the vaginal cell monolayer was studied to have a more comprehensive overview of the vaginal immune response in the presence of bacterial cells. This was achieved by performing the Olink® Target 96 assay (v.3026) (Olink® technology, Thermo Fisher Scientific), an immunoassay that measures a pattern of 92 inflammatory biomarkers, including inflammatory proteins and cytokines from various well-established families (full target list at https://olink.com/products-services/target/inflammation/). The assay measures each protein target individually, reporting relative levels as normalized protein expression (NPX) values on a log2 scale; thus, a change of 1 NPX unit reflects a twofold alteration in protein concentration. Briefly, after 24h incubation, supernatants were collected (centrifuged 2000xg for 10 min) and aliquoted (20 μL) into 96-well plates. Plates were stored at -20°C until Olink analysis. For each biomarker, the results were reported as a mean of NPX values within 3 replicates per condition. Biomarkers were included in the analysis if more than 55% of the replicates per condition had values above the limit of detection (LOD).

### 3D vaginal biomimetic model culturing

The 3D vaginal model was obtained by adapting the protocol from Edwards et al., 2022. Briefly, the bottom side of 12 well-plate thincerts (CELLSTAR® by Greiner Bio-One) was coated with 140 μL of rat tail collagen (Corning – Merck). The collagen solution was prepared by diluting 1:1.28 the rat tail collagen in 5X Roswell Park Memorial Institute RPMI:NaOH 1M (ratio 9:1) (Merck) and the pH adjusted at 6.5. Afterwards, the coated thincerts were let dry for 5h at room temperature followed by an incubation of 72h at 4°C. After 72h, BJ fibroblast cells (6x10^4^ cells/thincert) were added on top of the collagen layer and let them adhere for 6h, 37°C, 5% CO_2_. Inserts were then transferred to a deep 12-well plate (CELLSTAR® by Greiner Bio-One) adding BJ cells complete medium (EMEM) to the apical (200 μL) and the basolateral (5 mL) compartment and plates were returned to the incubator for 48h. Afterwards, vaginal epithelial cells (VK2/E6E7) (1x10^5^ cells/thincert) were seeded on the apical side and the plate was returned in the incubator for 48h. After refreshing the cell culture media on both the apical (KFSM complete medium) and basolateral sides (EMEM complete medium), the cells were incubated for two days before removing the KFSM medium from the apical side to establish an air-liquid interface. Afterwards, EMEM fresh medium was added only to the basolateral side every 3–4 days for 8 days. After 8 days, the histology of the 3D vaginal model was visualized via light microscopy. To do so, the apical and basolateral compartments were fixed in 4% paraformaldehyde (PFA). Transwell membranes were cut out of the insert and embedded in paraffin after which 5 µm transverse sections were stained with hematoxylin & eosin (HE). Imaging of the sections was performed on a Nikon Ti microscope.

### Bacterial adhesion and immune profiling in a 3D vaginal biomimetic model

To investigate bacterial interactions with the 3D vaginal biomimetic model, we assessed adhesion using two imaging techniques: confocal microscopy (CM) and scanning electron microscopy (SEM). For CM, bacteria were FITC stained beforehand as reported above. Afterwards, stained (CM) or unstained bacteria (SEM) suspensions (500 μL) were added to the apical compartment of the model (1x10^7^ - 1x10^8^ CFU), while the basolateral side was refreshed with 5mL EMEM medium, and incubated for 3h, at 37°C in 5% CO_2_. Adherent bacteria to the vaginal layers were washed twice in PBS and samples were treated differently depending on the technique used for imaging acquisition.

For CM, the apical and basolateral sides were fixed in 4% PFA for 30 min and kept overnight in PBS at 4°C. Afterwards, the vaginal cells nuclei were stained with Hoechst Nucleic acid (2µg/mL in PBS) (ThermoFisher Scientific) for 15 min at room temperature, at dark. After washing once in PBS, the vaginal cells plasma membranes were stained with deep red CellMask Plasma Membrane Stain (1µg/mL in PBS) (ThermoFisher Scientific) for 30 min at room temperature, at dark. After a wash in PBS, the transwells were incubated overnight 4°C in PBS for CM image acquisition. Imaging was performed using a Nikon Sora spinning disk confocal system mounted on a Nikon Ti-2 E inverted microscope. Excitation was achieved using lasers at 405 nm, 488 nm, and 640 nm. Fluorescence signals were detected with a Photometrics Kinetix camera. Image acquisition was controlled via Nikon NIS-Elements software. Images were captured using 40× objective lens (NA 0.95). Z-stacks were acquired with a 1 µm step size, and maximum intensity projections were generated for analysis.

For SEM, transwells containing 3D vaginal epithelial cells were fixed with 2.5% glutaraldehyde in 0.1M sodium cacodylate at 4°C. After rinsing with 0.1M sodium cacodylate containing 7.5% saccharose, samples were dehydrated in an ethanol series (50%-70%-90%-95%-100%) and hexamethyldisilizane (HMDS). Transwell membranes were cut out of the insert under a stereomicroscope, mounted on a stub with carbon tape and sputter coated with 10 nm gold. Imaging was performed using a JSM-IT100 (Jeol) at 10kV.

For immune assay, Olink® technology was used. Briefly, bacteria suspensions (1x10^6^ CFU) were added to the apical compartments for 24h, at 37°C in 5% CO_2_. Afterwards, supernatants from both apical and basolateral compartments were collected, centrifuged and aliquoted (20 μL) into 96-well plates. Plates were stored at -20°C until Olink® analysis and data were processed as reported above. Each condition was repeated 3 times.

### Analysis of immune biomarkers responses in human vaginal samples

Vaginal immune samples (n=185) from a clinical trial (NCT06425081) at the reference timepoint (healthy, baseline) were used. The protocol of this study was in accordance with the Declaration of Helsinski. The study and any amendments were approved by the ethical committee of the Antwerp University Hospital/University of Antwerp (Belgium, B3002024000026) and registered on clinicaltrials.gov (NCT06425081). Vaginal swabs were self-collected using a FLOQSwab® (Copan) and immediately transferred to a vial containing 1X Halt Protease Inhibitor Cocktail (Thermo Scientifix) and stored at 4°C until further processing. Samples were vortexed for 15-30 seconds and centrifuged, after which supernatant was stored at -80°C until further analysis. All samples were registered in the Biobank Antwerpen (Antwerp, Belgium; ID: BE 71030031000). Samples were analysed using the Olink® technology as described above. Every measurement above LOD was considered for analysis. Prevalence was calculated as the proportion of samples in which a marker was detected, divided by the total number of samples.

### Statistical analysis

Statistical analysis was performed using the software GraphPad Prism version 8.0.1 (GraphPad Prism Software Inc, San Diego, CA, USA, www.graphpad.com). Ordinary one-way/two-way ANOVA with Tukey correction was used for multiple comparisons. The Student’s t-test with Welch correction was used for two means comparison on a single variable. Differences were deemed significant for *p* < 0.05.

## Declaration statements Data Availability

The datasets supporting the findings of this study are available in the European open-access repository Zenodo (https://zenodo.org/) under the accession number **10.5281/zenodo.15388380.**

## Code availability

The analysis code supporting the findings of this study are available in the European open-access repository Zenodo (https://zenodo.org/) under the accession number **10.5281/zenodo.15388380.**

## Acknowledgments

Sincere thanks to the Laboratory of Applied Microbiology and Biotechnology team in Belgium, as well as the Beneficial Microbes lab team in Italy. The following colleagues have provided important support for the study, such as contributing to the Isala project management, strain collection and the logistics of lab, general genetic engineering advise and biosafety: Sarah Ahannach, Isabel Erreygers, Camille Gepts, Ines Tuyaerts, Nele Van Vliet and Max Dekeukeleire. All the icons and schematic images were created with BioRender.com (full license).

This work was supported by the European Research Council grant Lacto-Be (grant ID 852600) (awarded to SL and supports TE, JD) and proof-of-concept VALERIE (Horizon) (grant ID 101213306) (awarded to SL, supports JD) and ERC Runner-up ELLA (grant ID G0ATZ25N), FWO (G049022N, G031222N, SBO DeVeniR S006424N), the Inter-University Special Research Fund of Flanders (iBOF) for the POSSIBL project and the industrial research fund UAntwerpen for IOF POC project CRUCIAL. In addition, IS was supported by the University of Antwerp BOF-KP grant 53399, SW by a FWO postdoctoral grant 12AZ624N and CD by FWO doctoral grant 1S28622N. VC doctoral scholarship and part of the experimental activities were carried out in the Beneficial Microbes laboratory of the University of Bologna (Italy) and supported by the Italian Ministry of University and Research. The microscopes used in this publication, i.e. Tecnai Spirit G2 Biotwin (AUHA-08-004), Nikon SoRA (I003420N), were funded by Medium-scale research infrastructure grants of the FWO.

## Author contributions

Conceived and designed the experiments: VC, CD, TE, DV, CA, IP, ST, WM, MN, PAB, IS, CP, BV, SL

Performed the experiments: VC, CD, SB, IP, ST, WM, MN

Analyzed the data: VC, CD, TE, JD, TVR, DV, IP, ST, WM, MN, IS, CP, BV, SL

Contributed materials/analysis tools: VC, CD, TE, JD, TVR, EC, IVT, IP, ST, SW, WM, MN, PAB, IS, CP, BV, SL

Wrote the paper: VC, TE, TVR, IVT, IP, ST, WM, MN, PAB, SL

Revised the paper: VC, CD, TE, JD, TVR, EC, IVT, SB, DV, CA, IP, ST, WM, MN, PAB, SW, IS, CP, BV, SL

Funding acquisition and project management: BV, SL

