## Supplementary for "Exopolysaccharides of *Lactobacillus crispatus* mediate key balancing interactions with the vaginal mucosa"


### Table S1

| Species tested | Strains | Growth in MRS media ($OD_{600}$) Cut-off 1.5 | Growth in MRS+glycine2%+ sucrose2% ($OD_{600}$) Cut-off 1 | Chloramphenicol MIC (µg/mL) Cut-off 10 µg/mL |
|---|---|---|---|---|
| ***Lactobacillus crispatus*** | BC1 | 5.33 ± 1.01 | 1.53 ± 0.02 | 5 |
| | BC3 | 1.67 ± 0.12 | 1.25 ± 0.10 | 5 |
| | BC5 | 5.10 ± 0.52 | 1.64 ± 0.04 | 5 |
| | AMBV-0006 | 4.31 ± 1.14 | 1.28 ± 0.11 | 5 |
| | AMBV-0815 | 3.26 ± 0.09 | 1.64 ± 0.05 | 5 |
| | AMBV-1513 | 2.42 ± 0.67 | 1.14 ± 0.02 | 5 |
| | AMBV-2491 | 2.03 ± 0.88 | 1.44 ± 0.20 | 2.5 |
| ***Lactiplantibacillus plantarum*** | WCFS1 | 3.04 ± 0.32 | 1.50 ± 0.05 | 10 |

***Table S1. Selection of in-house L. crispatus strains stuitable for genetic manipulation. Data are reported as mean ± SD (n=2).*** *L. plantarum* WCFS1 was used as a control strain for electrocompetent cells preparation due to its high genetic accessibility using glycine-sucrose-based protocols.

### Figure S1

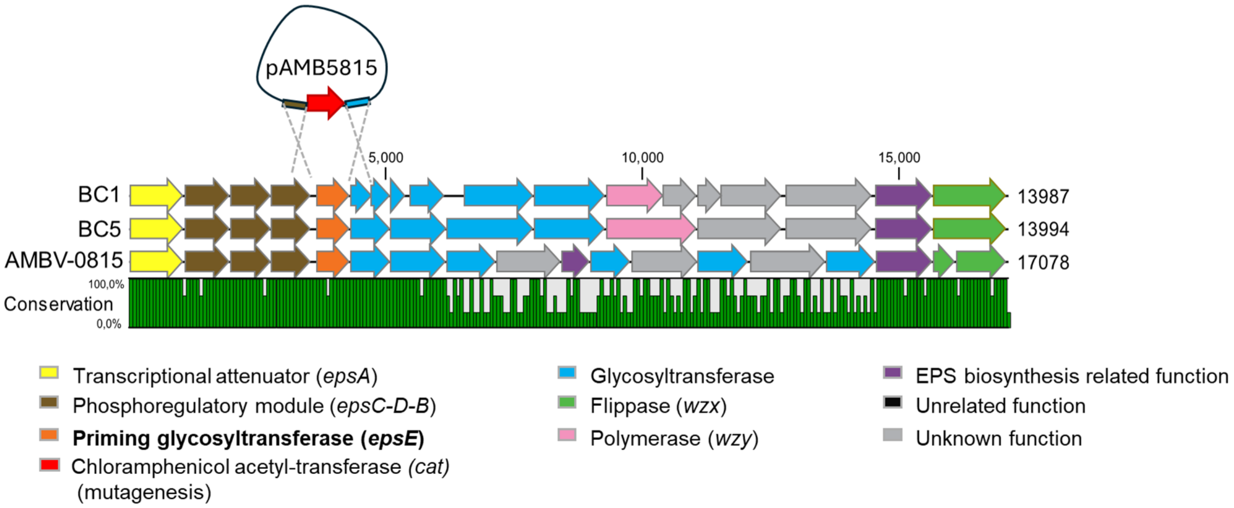


***Figure S1. Schematic representation of the EPS gene cluster in L. crispatus strains with highly conserved epsE neighboring genes and the construction of the semi-generic mutagenesis plasmid (pAMB5815).*** *Nucleotide sequence conservation between the strains is reported in the line plot (%).*

### Table S2

| **Species tested** | **Strains** | **Transformation efficiency (CFU/µg pNZ123)** | **Mutagenesis efficiency (pAMB5815) (CFU)** |
|---|---|---|---|
| ***Lactobacillus crispatus*** | BC1 | <$10^3$ | 0 |
| | BC5 | <$10^3$ | 0 |
| | AMBV-0815 | $1.5x10^4$ | 1 |
| ***Lactiplantibacillus plantarum*** | WCFS1 | $8.4x10^3$ | / |

***Table S2. Genetic accessibility of selected strains (pNZ123) and mutagenesis efficiency (pAMB5815).*** *L. plantarum* WCFS1 was used as a control strain for electrocompetent cells preparation due to its high genetic accessibility using glycine-sucrose-based protocols.

### Figure S2

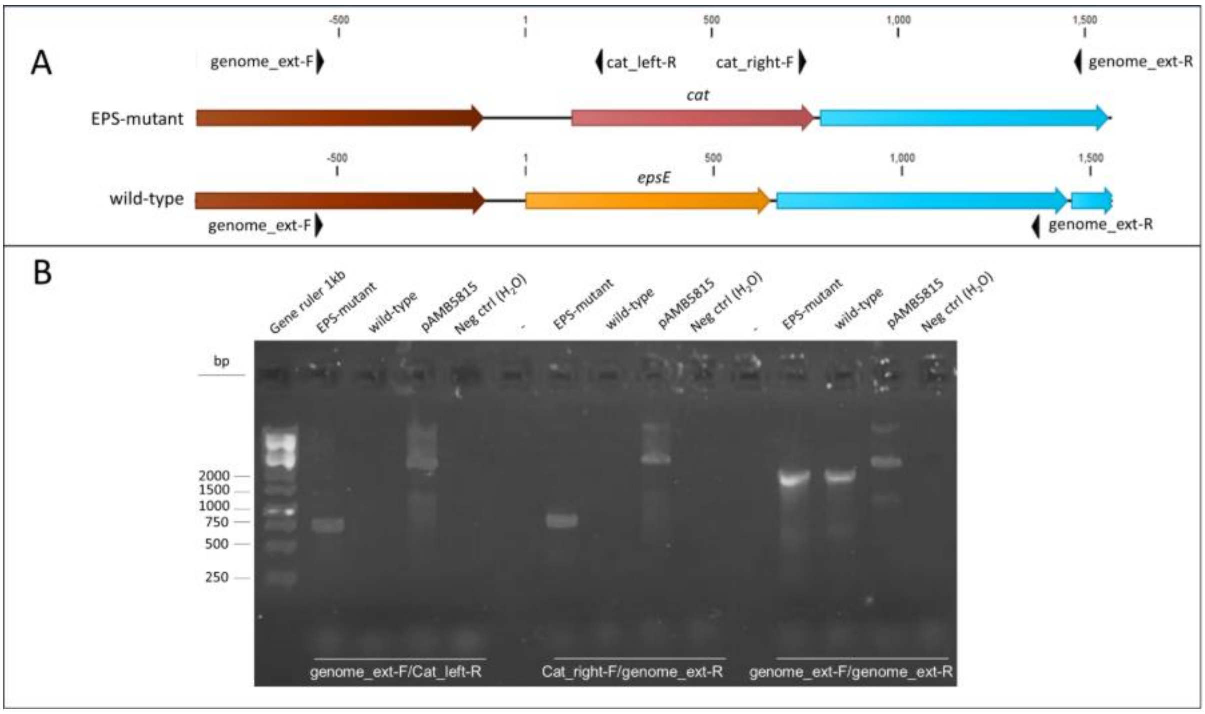


***Figure S2. A. Schematic representation of the primers used to check the double homologous recombination in L. crispatus AMBV-0815 ΔepsE (EPS-mutant). B. Gel electrophoresis (agarose 1% w/v) from colony PCR.*** *All samples were amplified with PCR using three different primer pairs, indicated in white. EPS-mutant and wild-type genome was obtained from colony PCR. Mutagenesis plasmid pAMB5815 and $H_2O$ were used as negative controls. Nucleotides position is reported starting from the start codon (ATG) of epsE gene in the wild-type strain. Fragments expected in EPS-mutant strain: genome_ext-F/Cat_left-R: 766 bp; Cat_right-F/genome_ext-R: 756 bp and genome_ext-F/genome_ext-R: 2048 bp. Fragments expected in wild-type strain: genome_ext-F/Cat_left-R: / ; Cat_right-F/genome_ext-R: / and genome_ext-F/genome_ext-R: 1920 bp. PCR amplicons obtained for EPS-mutant were Sanger sequenced.*

**Figure S3**

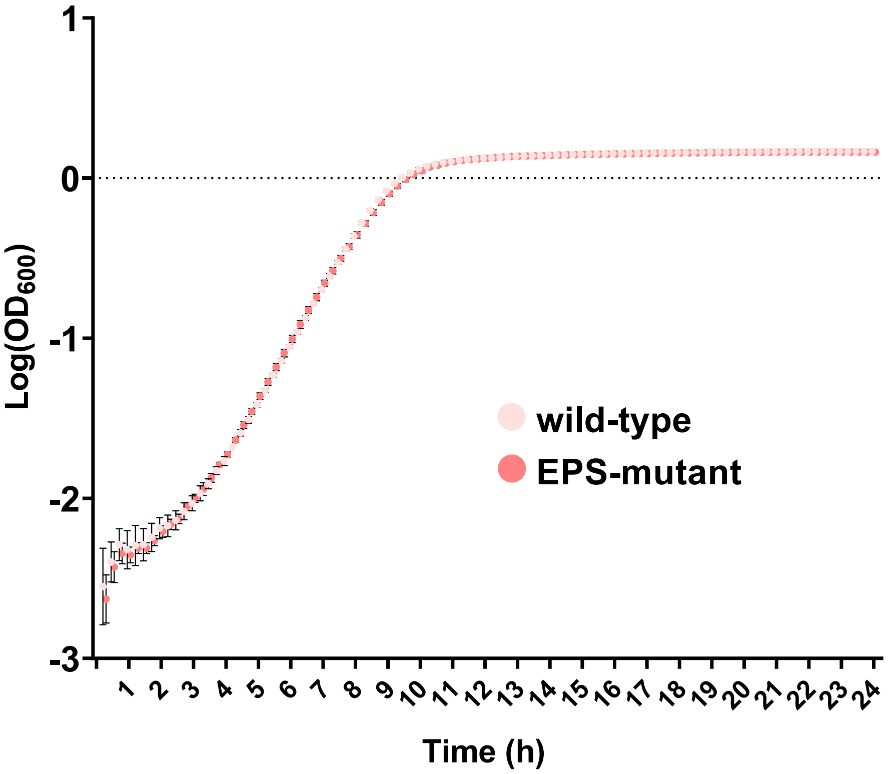


***Figure S3. Growth curve in MRS liquid of L. crispatus AMBV-0815 wild-type and EPS-mutant (24 h).*** *Data are reported as mean ± SD (n = 3).*

### Figure S4

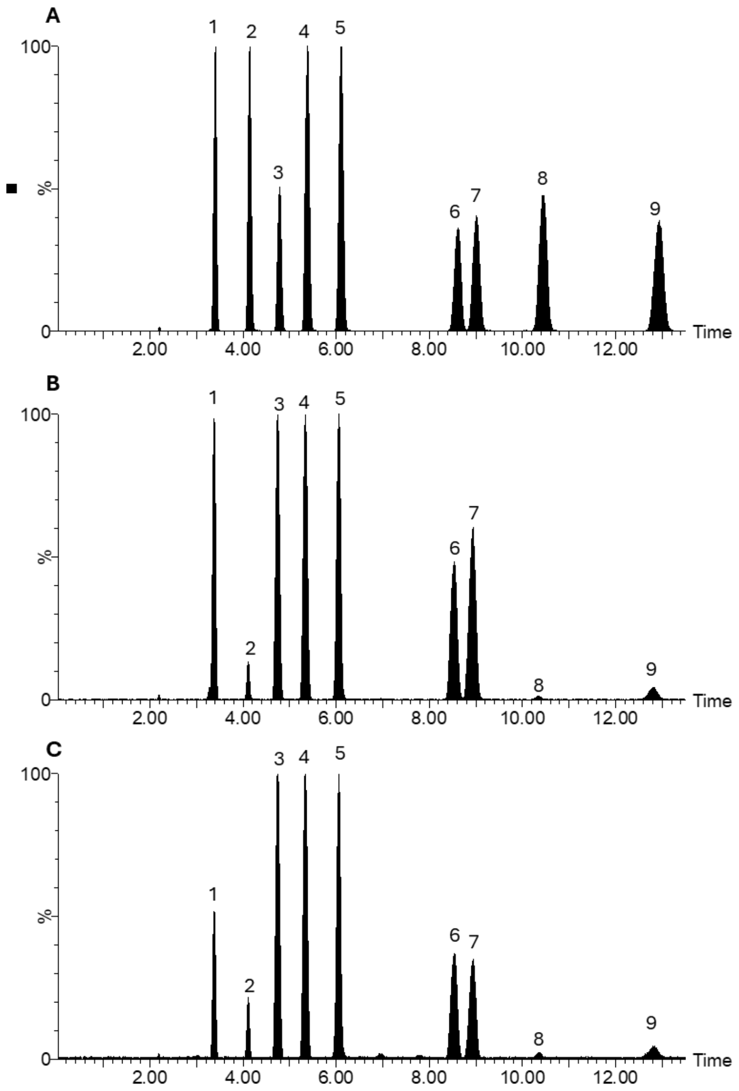


***Figure S4. Extract Ion Chromatograms of identified PMP-monosaccharides (LC-MS)****. (A) mixture of nine standard PMP-monosaccharides; (B) PMP-monosaccharides from L. crispatus AMBV-0815; (C) EPS-mutant samples. Peaks:* ***1****: D-Glucosamine;* ***2****: D-Mannose;* ***3****: Galactosamine;* ***4****: D-Ribose;* ***5****: D-Rhamnose;* ***6****: D-Glucose;* ***7****: D-Galactose;* ***8****: D-Xylose;* ***9****: D-Fucose.*

### Figure S5

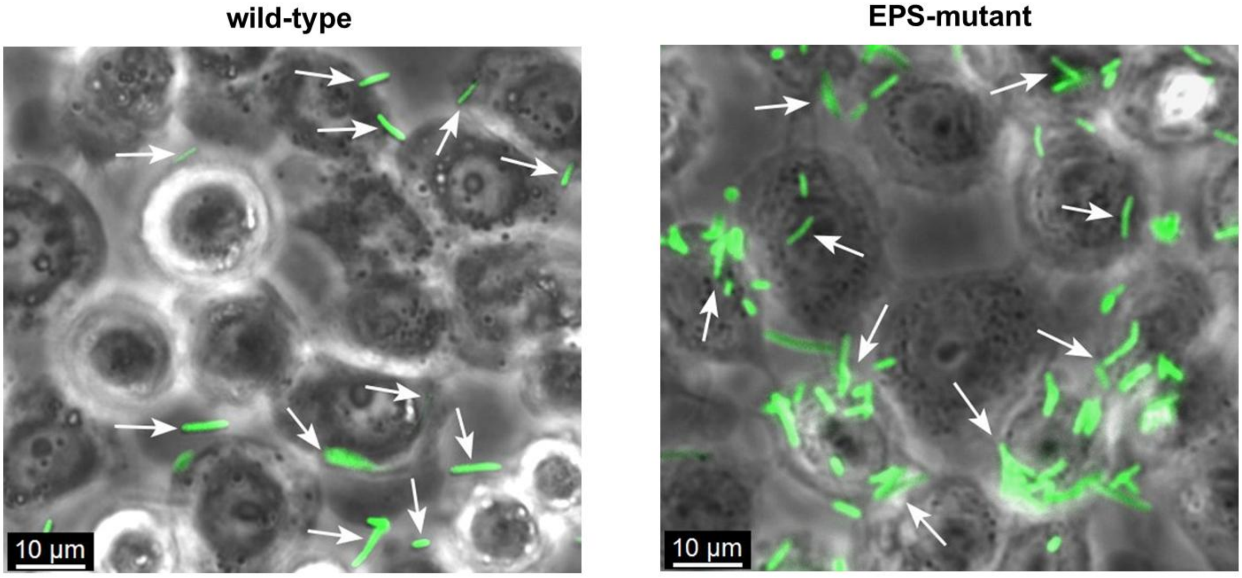


***Figure S5. Fluorescence microscopy micrographs of L. crispatus AMBV-0815 wild-type and EPS-mutant adherent on vaginal epithelium cells monolayer (VK2/E6E7) (3h).*** *L. crispatus AMBV-0815 wild-type and EPS-mutant were fluorescence stained with FITC (0.1 ug/mL - green channel, indicated with white arrows).*

### Figure S6

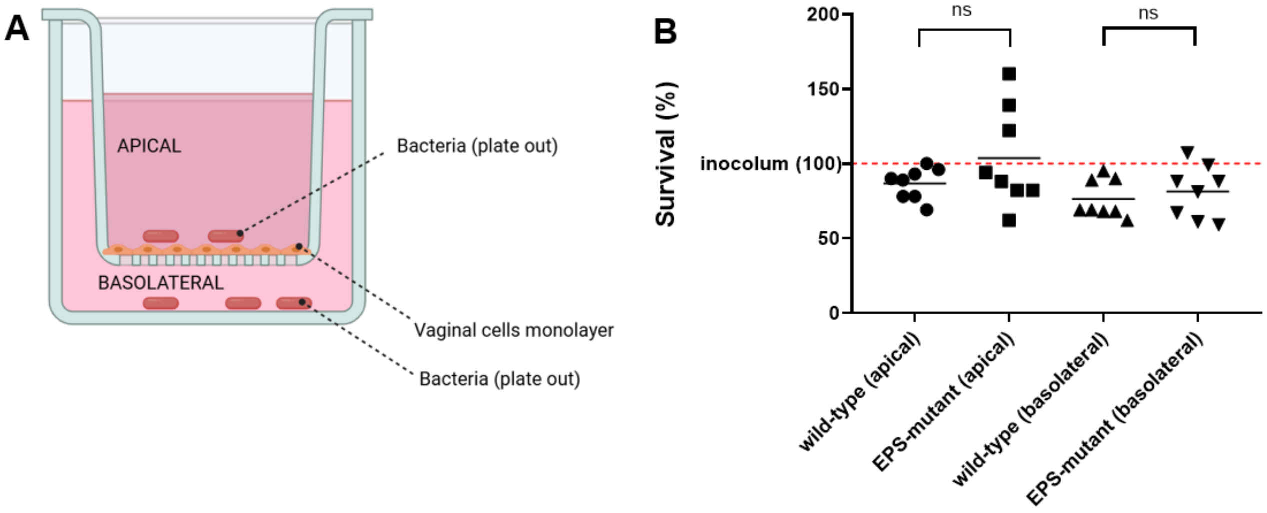


***Figure S6. Survival of bacterial cells in presence of vaginal cells on apical side (direct contact) and on basolateral side (indirect contact) of a transwell model (3h).*** *(A) Schematic representation of the assay. (B) Survival rates (%) of L. crispatus AMBV-0815 wild-type and EPS-mutant. The survival was calculated basing on the CFU plated out after 3h exposure in respect to the starting inoculum (CFU before exposure –100% survival). Two-way ANOVA with Tukey correction (n=8, ns =non significance, p > 0.05)*

### Figure S7

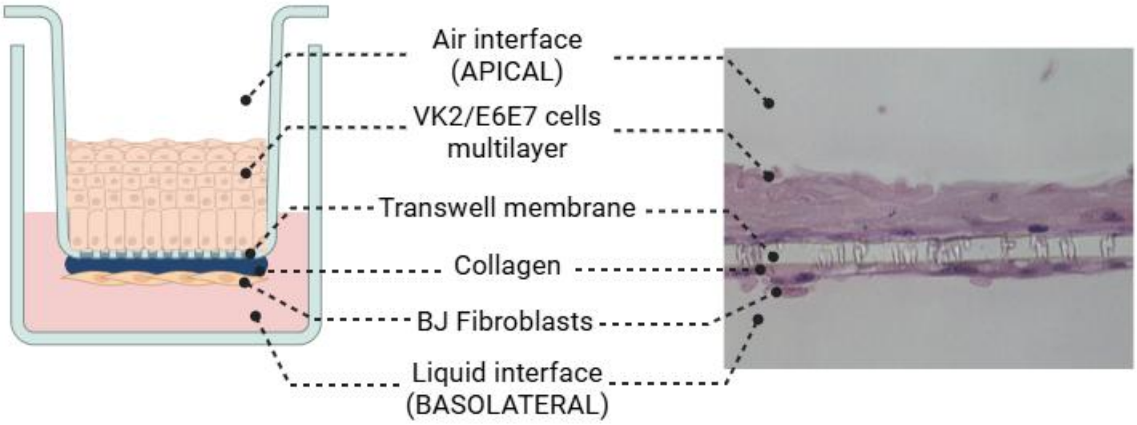


***Figure S7. Representation and micrograph of cross section of the 3D vaginal transwell model (hematoxylin and eosin stained) used in the present study.***
